# Hurricanes Pose Substantial Risk to New England Forest Carbon Stocks

**DOI:** 10.1101/2023.11.22.568264

**Authors:** Shersingh Joseph Tumber-Dávila, Taylor Lucey, Emery Boose, Danelle Laflower, Agustín León-Sáenz, Barry T. Wilson, Meghan Graham MacLean, Jonathan R. Thompson

**Author notes:** **Corresponding author**: Shersingh Joseph Tumber-Dávila.

## Abstract

Nature-based climate solutions are championed as a primary tool to mitigate climate change, especially in forested regions capable of storing and sequestering vast amounts of carbon. New England is one of the most heavily forested regions in the United States (>75% forested by land area), and forest carbon is a significant component of regional climate mitigation strategies. Large infrequent disturbances, such as hurricanes, are a major source of uncertainty and risk for policies that rely on forest carbon for climate mitigation, especially as climate change is projected to alter the intensity and geographic extent of hurricanes. To date, most research into disturbance impacts on forest carbon stocks has focused on fire. Here we show that a single hurricane in the region can down between 121-250 MMTCO_2_e or 4.6-9.4% of the total aboveground forest carbon, much greater than the carbon sequestered annually by New England’s forests (16 MMTCO_2_e yr^-1^). However, the emissions from the storms are not instantaneous; it takes approximately 19 years for the downed carbon to become a net emission, and 100 years for 90% of the downed carbon to be emitted. Using hurricane reconstructions across a range of historical and projected wind speeds, we find that an 8% and 16% increase in hurricane wind speeds leads to a 10.7 and 24.8 fold increase in the extent of high-severity damaged areas (widespread tree mortality). Increased wind speed also leads to unprecedented geographical shifts in damage; both inland and northward into heavily forested regions traditionally unaffected by hurricanes. Given that a single hurricane can emit the equivalent of 10+ years of carbon sequestered by forests in New England, the status of these forests as a durable carbon sink is uncertain. Understanding the risks to forest carbon stocks from large infrequent disturbances is necessary for decision-makers relying on forests as a nature-based climate solution.

## 1.0 Introduction

The impacts of climate change and the failure to meet emission reduction targets are driving a widespread interest in using nature-based climate solutions (NCS) to meet climate policy goals (Galik and Jackson 2009; Ellerman et al. 2016; Griscom et al. 2017; Roe et al. 2019). Forests are a major focus of NCS schemes, as they sequester the equivalent of nearly 25% of human carbon dioxide (CO2) emissions globally (Bonan 2008; Pan et al. 2011; Anderegg et al. 2022), with U.S. forests sequestering the equivalence of 10% of U.S. CO_2_ emissions (Birdsey et al. 2006). However, NCS policies often focus on the potential for future sequestration while ignoring the potential for existing carbon stocks to become a source of emissions due to disturbances (Anderson-Teixeira et al. 2013; Seidl et al. 2017; Brodribb et al. 2020). Therefore, relying on forest carbon offsets to mitigate greenhouse gas emissions has garnered considerable scrutiny, especially under the current regulatory and voluntary carbon market regimes (Haya 2010; Gifford 2020; Badgley et al. 2022a, b).

Using forests as NCS requires an accurate accounting of the risks posed by disturbance regimes, including climate stress, biotic agents, wildfires, and storms (i.e., snow, ice, lightning, and wind). The importance and complexity of accounting for these factors is magnified as climate change alters disturbance regimes and even introduces new confounding factors (Wu et al. 2023). For example, increased droughts will likely lead to increased susceptibility of trees to biotic agents, and an increased likelihood and magnitude of wildfires, especially in the Western U.S. (Anderegg et al. 2022). Under a changing climate, using historical data to calculate the likelihood of disturbance risks is likely inadequate; for example, the 100-year integrated risk of a moderate and severe wildfire across the U.S. has doubled from approximately 4% to 8% between the periods of 1984-2000 and 2001-2017 (Anderegg et al. 2020). Similarly, across the eastern U.S., warmer sea surface temperature will likely lead to more damaging tropical storms (Mann and Emanuel 2006).

Estimating impacts of disturbances on carbon stocks requires consideration of the disturbance properties, likely changes to those disturbances due to climate change, and the management response to the disturbance. Most previous research has focused on wildfires, which includes pyrogenic emissions from forest fires that are instantaneous (Campbell et al. 2007). In contrast, trees damaged by biotic agents and storms can either decompose in place over longer periods of time or potentially be stored in harvested wood products, all of which would uniquely affect the permanency of the forest carbon and may alter the balance between forests serving as a carbon source or sink (Fisk et al. 2013; Zscheischler et al. 2014; Seidl et al. 2017).

Throughout the last century, New England’s forests have served as a critical carbon sink, resulting from widespread reforestation following nineteenth century farm abandonment, reduced harvesting, and other land-use impacts (Albani et al. 2006; Bonan 2008). Currently, New England is among the most forested regions in the United States, with nearly 75% of its land covered by forests (Thompson et al. 2013; FIA USDA Forest Service 2022); sequestering 16 MMTCO_2_e of aboveground forest carbon annually (US EPA 2022). New England forest carbon is central to regional and national decarbonization strategies, as many states strive to become “Net-Zero’’ emitters in the coming decades (Wayburn 2009; US Climate Change Science Program 2014; Thompson et al. 2020; Higgins et al. 2021), and as industries begin to take part in forest offset markets (Kerchner and Keeton 2015).

Hurricanes are a dominant disturbance agent in New England, with the North Atlantic Basin being amongst the most active regions for tropical cyclones, resulting in New England being impacted by catastrophic hurricanes about once a century (Boose et al. 2001; Landsea et al. 2015). Ten hurricanes had a significant impact in New England during the 20th century, the most impactful being: the Great New England Hurricane of 1938, Carol in 1954, and Bob in 1991 (Landsea et al. 2015). For example, the 1938 hurricane downed 70% of the timber volume at Harvard Forest in central Massachusetts (Foster and Boose 1992). It caused extensive damage throughout New England, destroying over 8,900 buildings and damaging an additional 15,000 (Massachusetts Office of Coastal Zone Management 2002; Long 2016). It has been suggested that over the next century, storm wind speeds may increase by 6-16% due to increases in Atlantic basin sea surface temperatures (Emanuel 2005; Knutson et al. 2009; Bender et al. 2010). It is unknown whether the frequency of storms will change; however, some meteorologists predict that climate change may lead to fewer, yet more intense hurricanes, with the probability of storm impacts to increase by 200-300% throughout the next century (Emanuel 2005; Mann and Emanuel 2006; Knutson et al. 2009; Bender et al. 2010).

In this study, we quantify the potential impact of 21st century hurricanes on New England forest aboveground forest carbon stocks. We analyze three scenarios based on hurricane data for the ten most impactful storms in the 20th century: 1) Baseline— no change in hurricane wind intensity; 2) Projected—8% increase in wind speeds from the baseline; and 3) Maximum Severity—16% increase in wind speeds. The extent and intensity of the modeled storms, together with a novel map of forest composition and carbon density, and a harvested wood products model are used to estimate the impact that storms would have on aboveground forest carbon. Specifically, we ask: 1) What risks do hurricanes pose to existing live aboveground forest carbon stocks in New England? 2) How will this risk be affected by predicted changes in climate and subsequent altered wind disturbance regimes affecting the intensity and geographic impact of hurricanes?

## 2.0 Materials and methods

### 2.1 Estimating Hurricane Impacts on Aboveground Forest Carbon

Our aim is to estimate the forest carbon losses that would occur during a hurricane in New England. The four major components needed to make this estimation are: 1. spatial reconstruction of hurricane paths and their Enhanced Fujita (EF) damage (see methods section 2.2), 2. maps of aboveground forest carbon for each of eight tree type-height vulnerability classes (2.4), 3. an estimate of the expected percent of trees downed by each experienced EF damage for each tree vulnerability class (2.3), and 4. a harvested wood products model to estimate the carbon emissions pathways (2.5; Figure 1). We combined the first three components to calculate the amount of forest carbon downed within each forested pixel in New England based on the EF rating and the tree vulnerability classification. We did this for all 10 storms in each of the three scenarios (30 storms total). We then calculate the amount of downed forest carbon within each state and county following each storm, as well as the size and strength of each hurricane. Finally, we estimated the carbon emissions from downed forest carbon post-hurricane using a harvested wood products carbon storage and emissions model.

**Figure 1.**
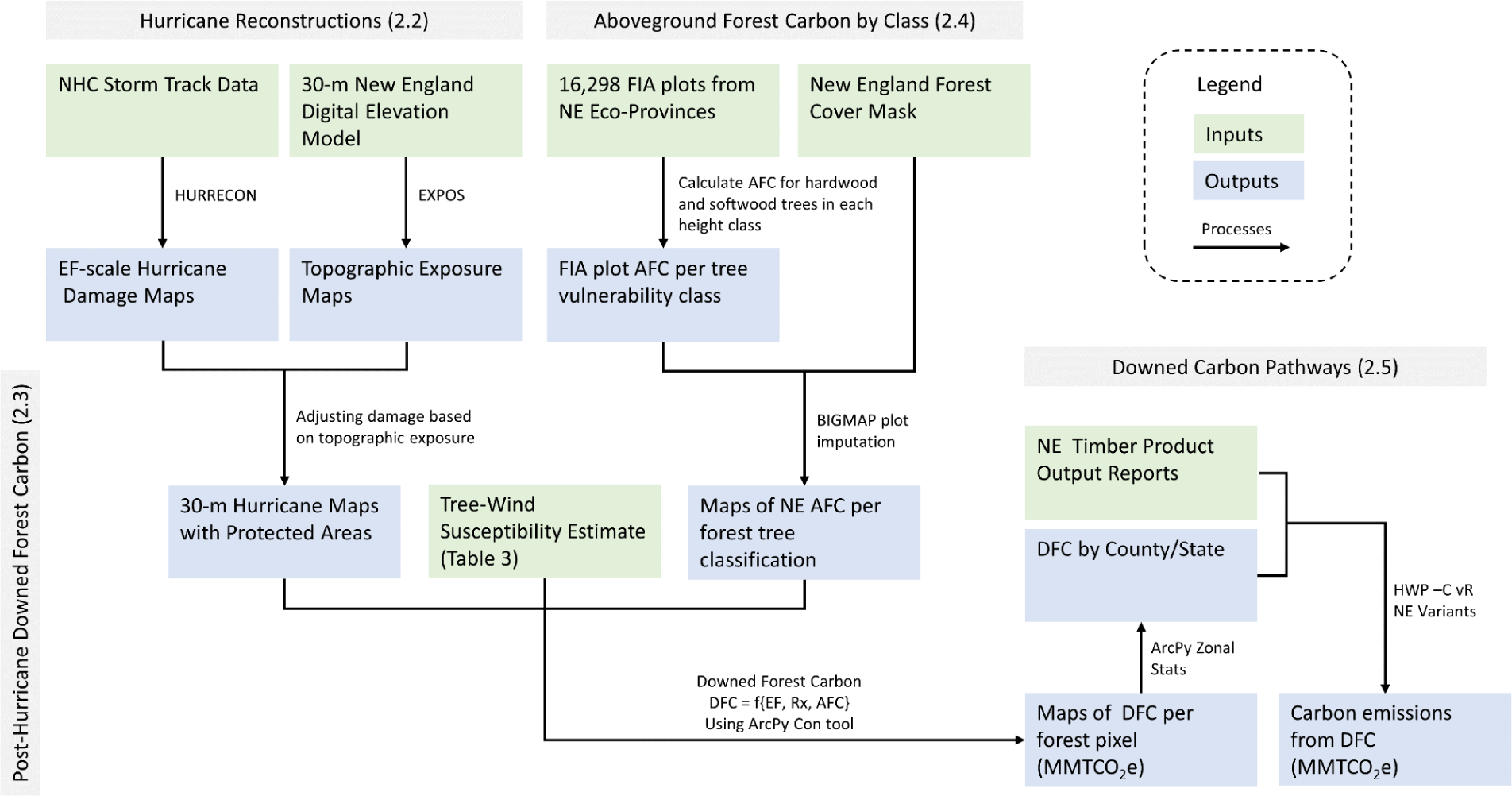
Flowchart of methods to calculate downed forest and the emissions from downed forest carbon by combining the hurricane reconstructions (2.2) with the New England aboveground forest carbon estimations (2.4), calculated using the forest tree vulnerability to hurricane-force winds (2.3), followed by the calculations of emissions using the harvested wood products model (2.5). Inputs are represented by green boxes, outputs by blue boxes, and processes (models and major analyses) by arrows. The gray headers represent the different major processes, described in the methods subsections in parentheses.

### 2.2 Hurricane Reconstructions & Scenarios

We modeled the impacts of ten 20^th^ century hurricanes that caused EF1 or higher damage in New England (Table 2 & Figure 2). These hurricanes were chosen because of the abundance of meteorological and damage data for these storms and because the 20^th^ century is reasonably typical of the 400-year period since European settlement, with somewhat less hurricane activity than the 19^th^ century and somewhat more than the 18^th^ century (Boose et al. 2001). Each storm was modeled as if it occurred in 2020, and each storm was simulated under three disturbance regime scenarios: (1) baseline—actual historical wind speeds from HURDAT2 (Landsea et al. 2015), (2) projected—wind speeds increased by 8%, and (3) maximum severity—wind speeds increased by 16% (Figure S2).

**Figure 2.**
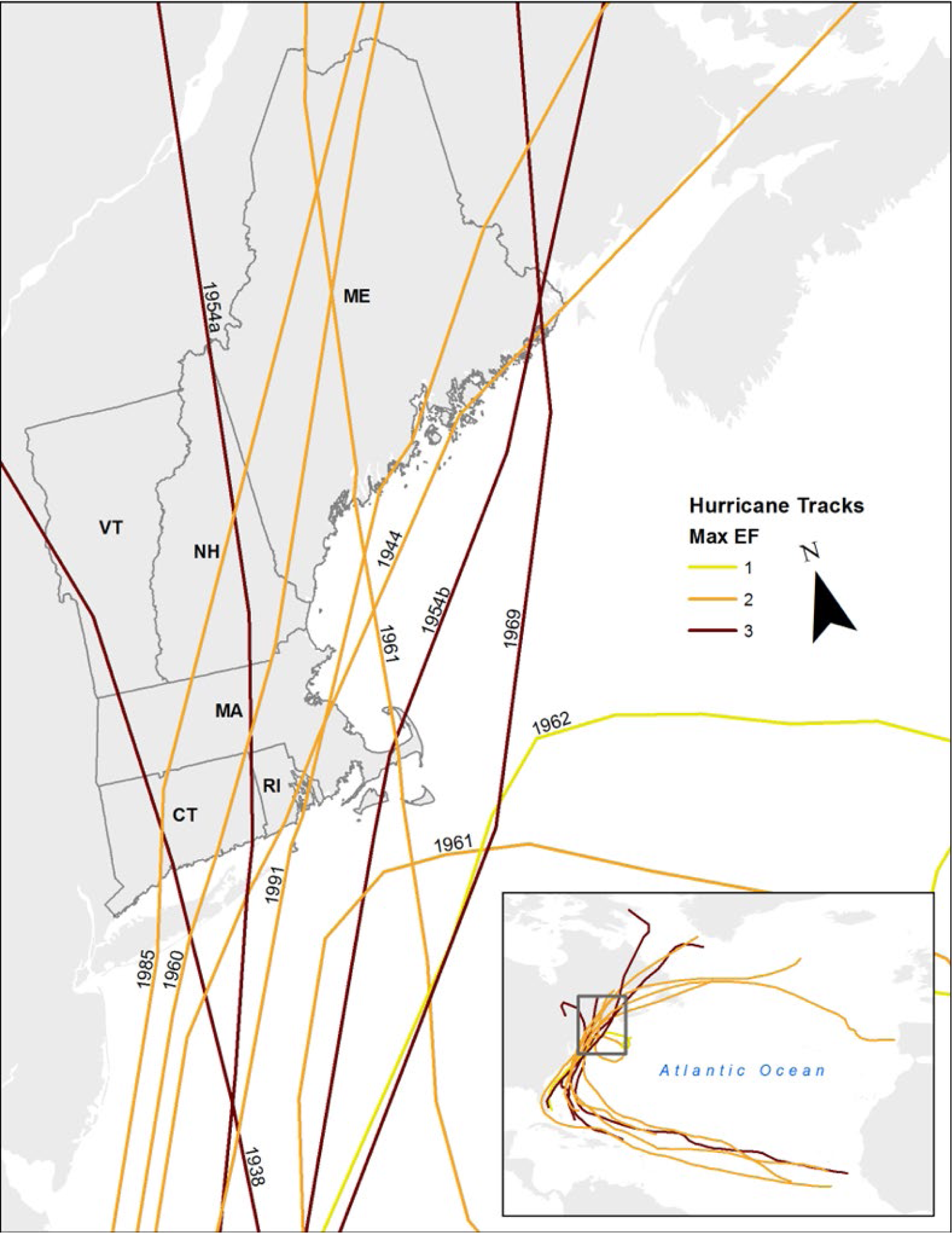
Tracks for the ten most impactful 20th century New England hurricanes. The colors indicate the maximum EF value to impact New England (generally the location where the storm made landfall in the region, as storms weaken throughout their trajectory). The EF values represent the baseline scenario, which is the historical strength of the hurricane on record. The inset map on the bottom right shows the entire hurricane tracks across the Atlantic basin with the gray box depicting the region of the main map.

**Table 1.**
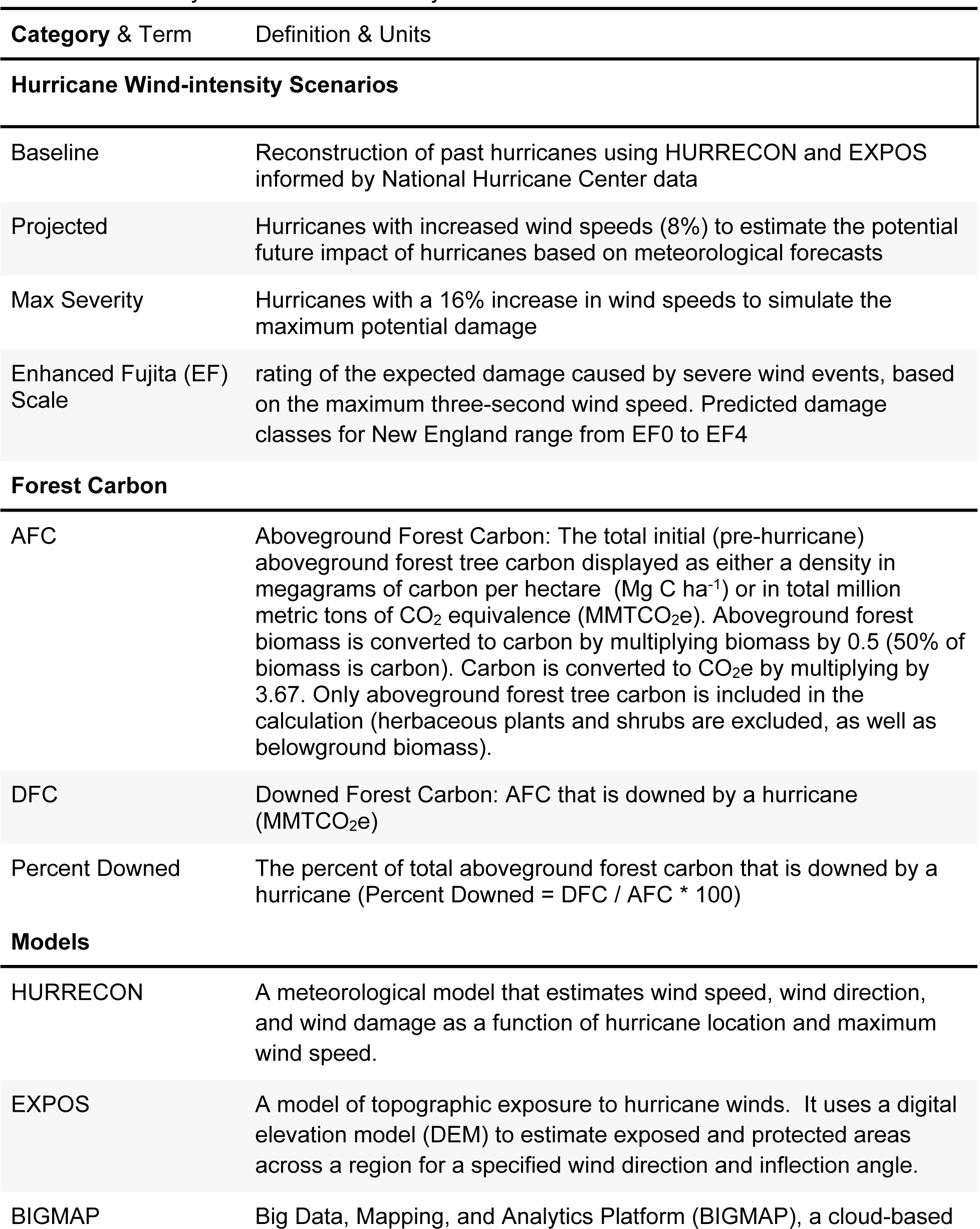
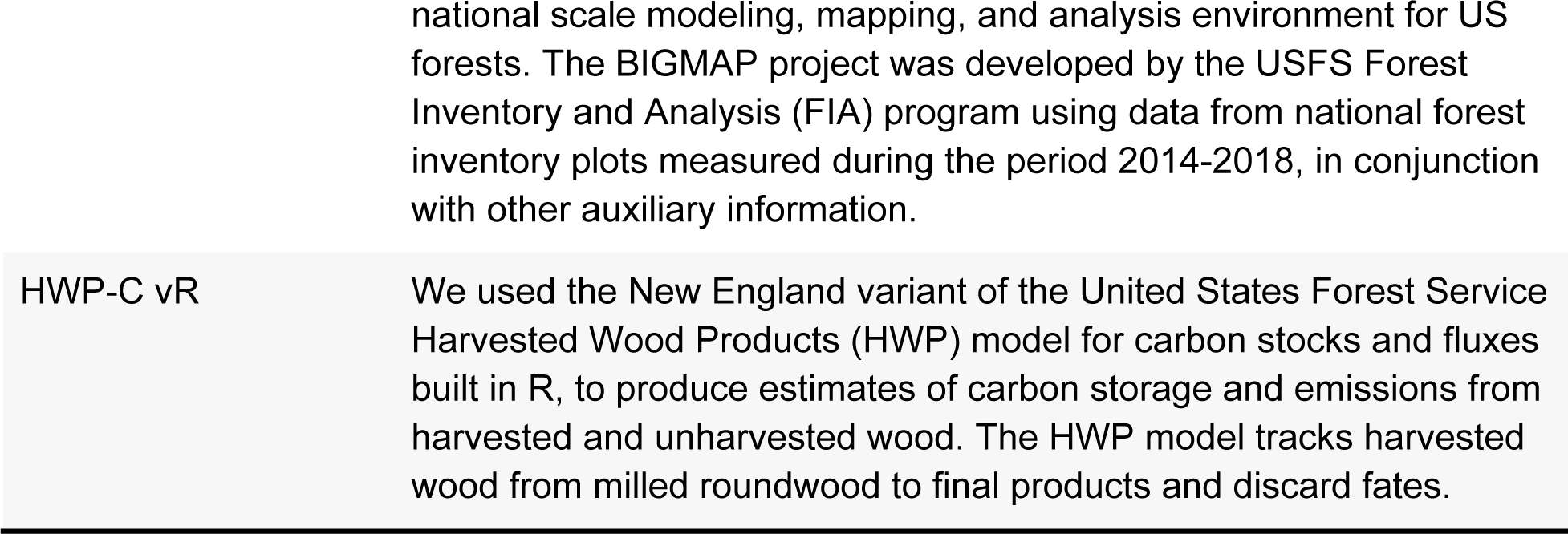
Commonly used terms and acronyms.

**Table 2.** Ten 20th century hurricanes and their impact in terms of affected area and downed forest carbon across Enhanced Fujita (EF) classes for each of the three hurricane intensity scenarios.

|  | Total Impacted Area |  | Damaged Area by EF Class (km<sup>2</sup>) |  |  |  | Downed Forest Carbon |  |
| --- | --- | --- | --- | --- | --- | --- | --- | --- |
|  | km<sup>2</sup> | % Area | EF0 | EF1 | EF2 | EF3 | MMTCO<sub>2</sub>e | % Down |
| **Great New England (1938)** |  |  |  |  |  |  |  |  |
| Baseline | 68,666 | 40.2% | 27,648 | 23,460 | 16,543 | 1,015 | 280 | 10.5% |
| Projected | 90,619 | 53.0% | 41,882 | 19,756 | 22,684 | 6,298 | 389 | 14.6% |
| Max Severity | 108,713 | 63.6% | 52,633 | 20,765 | 21,820 | 13,494 | 472 | 17.7% |
| **Great Atlantic (1944)** |  |  |  |  |  |  |  |  |
| Baseline | 45,905 | 26.9% | 22,413 | 22,916 | 576 | 0 | 126 | 4.7% |
| Projected | 55,997 | 32.8% | 24,074 | 26,193 | 5,730 | 0 | 179 | 6.7% |
| Max Severity | 72,395 | 42.4% | 33,723 | 23,126 | 15,546 | 0 | 259 | 9.7% |
| **Carol (1954)** |  |  |  |  |  |  |  |  |
| Baseline | 76,389 | 44.7% | 30,596 | 31,289 | 14,503 | 0 | 270 | 10.2% |
| Projected | 95,224 | 55.7% | 39,845 | 28,249 | 22,596 | 4,534 | 373 | 14.0% |
| Max Severity | 112,681 | 65.9% | 48,160 | 26,337 | 26,984 | 11,200 | 481 | 18.1% |
| **Edna (1954)** |  |  |  |  |  |  |  |  |
| Baseline | 38,967 | 22.8% | 31,628 | 7,071 | 267 | 0 | 81 | 3.1% |
| Projected | 64,955 | 38.0% | 49,234 | 14,527 | 1,194 | 0 | 154 | 5.8% |
| Max Severity | 85,742 | 50.2% | 58,707 | 23,699 | 3,187 | 149 | 222 | 8.3% |
| **Donna (1960)** |  |  |  |  |  |  |  |  |
| Baseline | 32,218 | 18.8% | 21,405 | 10,813 | 0 | 0 | 69 | 2.6% |
| Projected | 43,186 | 25.3% | 25,478 | 17,101 | 607 | 0 | 109 | 4.1% |
| Max Severity | 52,755 | 30.9% | 26,165 | 21,527 | 5,063 | 0 | 160 | 6.0% |
| **Esther (1961)** |  |  |  |  |  |  |  |  |
| Baseline | 31,824 | 18.6% | 25,266 | 6,558 | 0 | 0 | 65 | 2.4% |
| Projected | 41,391 | 24.2% | 28,982 | 12,409 | 0 | 0 | 101 | 3.8% |
| Max Severity | 50,493 | 29.6% | 31,124 | 18,315 | 1,054 | 0 | 133 | 5.0% |
| **Alma (1962)** |  |  |  |  |  |  |  |  |
| Baseline | 6,578 | 3.8% | 6,439 | 139 | 0 | 0 | 9 | 0.3% |
| Projected | 13,239 | 7.8% | 12,034 | 1,205 | 0 | 0 | 20 | 0.8% |
| Max Severity | 23,255 | 13.6% | 20,735 | 2,520 | 0 | 0 | 41 | 1.5% |
| **Gerda (1969)** |  |  |  |  |  |  |  |  |
| Baseline | 20,468 | 12.0% | 18,929 | 1,539 | 0 | 0 | 32 | 1.2% |
| Projected | 50,298 | 29.4% | 45,616 | 4,610 | 71 | 0 | 98 | 3.7% |
| Max Severity | 80,145 | 46.9% | 68,139 | 10,925 | 1,080 | 0 | 177 | 6.6% |
| **Gloria (1985)** |  |  |  |  |  |  |  |  |
| Baseline | 61,627 | 36.1% | 38,935 | 22,692 | 0 | 0 | 148 | 5.6% |
| Projected | 76,442 | 44.7% | 43,213 | 31,311 | 1,919 | 0 | 206 | 7.7% |
| Max Severity | 89,521 | 52.4% | 43,139 | 36,712 | 9,670 | 0 | 278 | 10.5% |

| **Bob (1991)** |  |  |  |  |  |  |  |  |
| --- | --- | --- | --- | --- | --- | --- | --- | --- |
| Baseline | 51,350 | 30.0% | 30,447 | 19,553 | 1,349 | 0 | 135 | 5.1% |
| Projected | 65,780 | 38.5% | 34,522 | 24,880 | 6,378 | 0 | 196 | 7.4% |
| Max Severity | 82,572 | 48.3% | 41,621 | 26,630 | 14,028 | 293 | 279 | 10.5% |
| **Storm Averages** |  |  |  |  |  |  |  |  |
| Baseline | 43,399 | 25.4% | 25,371 | 14,603 | 3,324 | 102 | 121 | 4.6% |
| Projected | 59,713 | 34.9% | 34,488 | 18,024 | 6,118 | 1,083 | 182 | 6.9% |
| Max Severity | 75,827 | 44.4% | 42,415 | 21,056 | 9,843 | 2,514 | 250 | 9.4% |

The HURRECON model is a simple meteorological model that estimates wind speed, wind direction, and wind damage as a function of hurricane location and maximum wind speed. The model is based on empirical studies of many hurricanes and can generate results for a single site or an entire region. The updated version of HURRECON used in this study (Boose 2023a) uses the same equations to estimate wind speed and direction as the original model (Boose et al. 2001). New features include the ability to estimate wind damage on the enhanced Fujita scale (Edwards et al. 2013) instead of the older Fujita scale (Fujita 1971) and imports hurricane track and intensity data directly from the U.S. National Hurricane Center’s HURDAT2 database (Landsea et al. 2015). The enhanced Fujita scale is used rather than the more common Saffir-Simpson scale because the former characterizes wind damage at a specific location, while the latter characterizes maximum wind damage anywhere in a hurricane.

Output from HURRECON informs the EXPOS model, which is a simple model of topographic exposure to hurricane winds. It uses a digital elevation model (DEM) to estimate exposed and protected areas at the pixel level, for a specified wind direction and inflection angle. The revised version of the model used in this study (Boose 2023b) uses the same algorithms to calculate exposure as the original model (Boose et al. 2001). New features include the ability to refine regional maps of wind damage from HURRECON by reducing the level of predicted wind damage at locations that are topographically protected from the predicted peak wind direction at that location. For this study we used a 30-meter digital elevation model for New England from the Shuttle Radar Topography Mission (USGS EROS 2018) and a 6-degree inflection angle informed by previous EXPOS studies (Boose et al. 1994, 2001, 2004). Damage in protected areas was reduced by two enhanced Fujita classes (e.g., a protected pixel with an EF2 rating was downgraded to EF0).

### 2.3 Forest Tree Vulnerability to Hurricane-force Winds

Based on the EF-scale damage predicted for a given location in New England, and the forest composition at the time of the hurricane, we estimate the degree of damage and translate that to percent tree mortality and forest carbon loss. To predict the impact that hurricanes have on aboveground forest carbon, we prescribed the probability that a forest tree is downed by a hurricane based on the two major axes of tree susceptibility to windthrow: 1) tree height, and 2) hardwood vs. softwood (Raymer 1962; Cooper-Ellis et al. 1999; Canham et al. 2001; Busing et al. 2009). The probability of downed trees (Table 3) following hurricane disturbances is informed by 1) the observed tree mortality of various species following past wind disturbances (Foster 1988; Godfrey and Peterson 2017), 2) differences in the plant structural traits of the common hardwood and softwood species in New England, with conifers tending to be more susceptible to hurricane-force winds (Cooper-Ellis et al. 1999; Busing et al. 2009), and 3) the observation that taller trees are more susceptible to windthrow (Raymer 1962; Canham et al. 2001). Under our framework, tall conifers are most susceptible to windthrow, and short hardwoods are least susceptible (Table 3). It is important to note that we only investigated the impacts from hurricane-force winds, and not those of storm-surge and/or precipitation that can also lead to catastrophic damages, especially along the coast and steep hillslopes.

**Table 3.** Forest tree vulnerability to hurricane-force winds by tree type and height across the Enhanced Fujita Scale classes.

| Enhanced<br>Fujita<br>Scale | Max 3-second<br>Wind<br>Gust<br>(mph) | Expected Tree<br>Damage* | Down Probability (%) by Tree Height<br>(Hardwood - Softwood) |  |  |  |
| --- | --- | --- | --- | --- | --- | --- |
|  |  |  | 0-10 m | 10-20 m | 20-30 m | 30+ m |
| EF0 | 65-85 | Leaves and fruit<br>off, branches<br>broken, tree<br>damage | 5-10% | 10-15% | 15-20% | 20-25% |
| EF1 | 86-110 | Trees blown down | 10-15% | 15-25% | 25-35% | 35-45% |
| EF2 | 111-135 | Extensive blow<br>downs | 15-25% | 35-50% | 55-75% | 70-95% |
| EF3 | 136-165 | Most trees down | 25-35% | 45-60% | 65-85% | 85-100% |
| EF4 | 166-200 | Catastrophic forest<br>damage | 95% | 100% | 100% | 100% |

\*Adapted from Table 1 of Boose et al. 2001

### 2.4 Mapping New England’s Forested Landscape

We mapped aboveground forest carbon for each of the 8 tree susceptibility categories representing hardwood and softwood trees across the range of tree heights (Table 3). We used the Big Data, Mapping, and Analytics Platform (BIGMAP), a cloud-based national scale modeling, mapping, and analysis environment for US forests. The BIGMAP project was developed by the USFS Forest Inventory and Analysis (FIA) program using data from national forest inventory plots measured during the period 2014-2018, in conjunction with other auxiliary information. Vegetation phenology derived via harmonic regression of Landsat 8 OLI scenes collected during the same time period, along with climatic and topographic raster data, were processed to create an ecological ordination model of tree species and produce a feature space of ecological gradients that was used to impute FIA plot data to pixels, and assign values for key forest inventory variables (Ohmann and Gregory 2002; Wilson et al. 2012, 2013, 2018). For our study, the key variable was live aboveground forest carbon across the eight tree-species-height categories (Table 3).

To create our desired BIGMAP product, we gathered data from 16,298 national forest inventory plots (measured between 2014-2018) from across the three ecosystem provinces that are represented by New England forests (212-Laurentian Mixed Forest Province, M212-Adirondack-New England Mixed Forest, and 221-Eastern Broadleaf Forest). For each plot, we used the FIA tree table and inferred tree heights when necessary, using the appropriate site index curve equations (Carmean et al. 1989). We then calculated the aboveground tree carbon across the eight tree height and hardwood/softwood classes for each inventory plot. These data were extracted from the BIGMAP plot imputation model and resulted in eight 30-meter resolution raster products of predicted aboveground forest carbon for each of the tree susceptibility categories across New England Forests.

### 2.5 Harvested Wood Products Carbon Storage & Emissions Estimates

We used the New England variant of the state level Harvested Wood Products (HWP) model, HWP-C vR (based on the national level model, USFS HWP-C v1; Anderson et al. 2013) to produce estimates of carbon storage and emissions from harvested and unharvested wood. The HWP model tracks harvested wood from milled roundwood to final products and discard fates. HWP-C vR has been used for California, Oregon, and Washington wood products carbon inventories (Groom and Tase 2022; Lucey et al. 2023), and we parameterized this model for New England. Carbon storage pool estimates include products in use (PIU), solid waste disposal sites such as dumps and landfills (SWDS), and remaining downed wood from storm events (DFC_s_). Downed wood from storm events also includes any biomass left in the forest after salvage harvest, and the decay of the downed wood (DFC) was modeled using species group average decay rates (Russell et al. 2014). Carbon emissions pools include emitted with energy capture (i.e., fuelwood or burned onsite at mills for energy; EEC), emitted without energy capture (e.g., decay from SWDS; EWoEC), and decay from downed wood left in the forest (DFC_e_). Estimated pools and emissions only represent carbon from trees damaged by the storm. Carbon sequestration from regeneration post-hurricane disturbance was not included in these pools or in our analyses.

Using the most recent New England Timber Products Output reports (FIA USDA Forest Service 2018), we calculated proportions of logging residues by species (reflects harvesting efficiencies), mill residues, and timber product ratios separately for northern and southern New England counties. The northern New England variant included three counties in New Hampshire, five counties in Vermont, and all of Maine. Southern New England included the remaining counties and states in New England. Primary product ratios for New England were created using the most recent northeast regional Timber Product Output report, and national end use ratios (McKeever 2009; McKeever and Howard 2011) were used to estimate proportions of biomass going to end uses as well as decomposition rates after wood products were discarded. From these ratios, we estimate a small proportion of harvested wood is manufactured into short-lived products or emitted during the milling process. The remaining primary products are turned into short- and longer-lived products based on species group, timber product, and most common end use products for these groups (Groom and Tase 2022).

Former variants of the HWP carbon model were intended to estimate cumulative carbon storage for products in use and in solid waste disposal sites over time. These estimates were created using annual historical harvest volume records. There were no historical harvest volume records or simulated future harvest volume used for this analysis, only the salvaged wood following the simulated hurricane disturbance. Therefore, we used the HWP model to estimate only the fate of carbon stored and emitted from the salvage harvest following each storm event. We assumed that on average, 25% of down wood would be salvage-harvested after each storm and that salvage harvest occurred the same year as the storm. The ratio of salvage harvest is based on historical salvage rates following hurricane disturbances affecting forested regions (Foster et al. 1997; Foster and Orwig 2006; Stanturf et al. 2007), and is limited by sawmill, storage, and transportation capacities, as well economic pressures (Sanginés de Cárcer et al. 2021). Exact salvage rates and timber product ratios were based on size criteria (height) and hardwood or softwood species. Approximately 26% of the largest size class trees (21m +) were removed for use in sawtimber and 10% of the medium sized trees (between 11m - 20m in height) were also removed for poletimber, to simulate targeted salvage logging of the most usable wood. We ran the HWP model with salvage harvest volume for each of the ten simulated storms at all three wind intensity scenarios. We then averaged the outputs by county within each wind intensity scenario.

## 3.0 RESULTS

### 3.1 Current status of New England Forest Carbon

New England is 75% forested by area, with an average AFC density of 56.3 Mg ha^-1^ for forested areas (Table 4), according to our BIGMAP product. Rhode Island is the least forested state, 51% of total land area, while Maine contains 55% of all New England’s forests, but has the lowest AFC density at 45.8 Mg ha^-1^ (Table 4). Connecticut and Massachusetts have the highest AFC densities, 70.1 and 70.8 Mg ha^-1^ respectively, but are only 56% forested, while New Hampshire and Vermont are both similarly densely forested (∼68 Mg ha^-1^ AFC) and have high forest cover (80% and 74% forested by area; Table 4; Figures 3 & S1).

**Figure 3.**
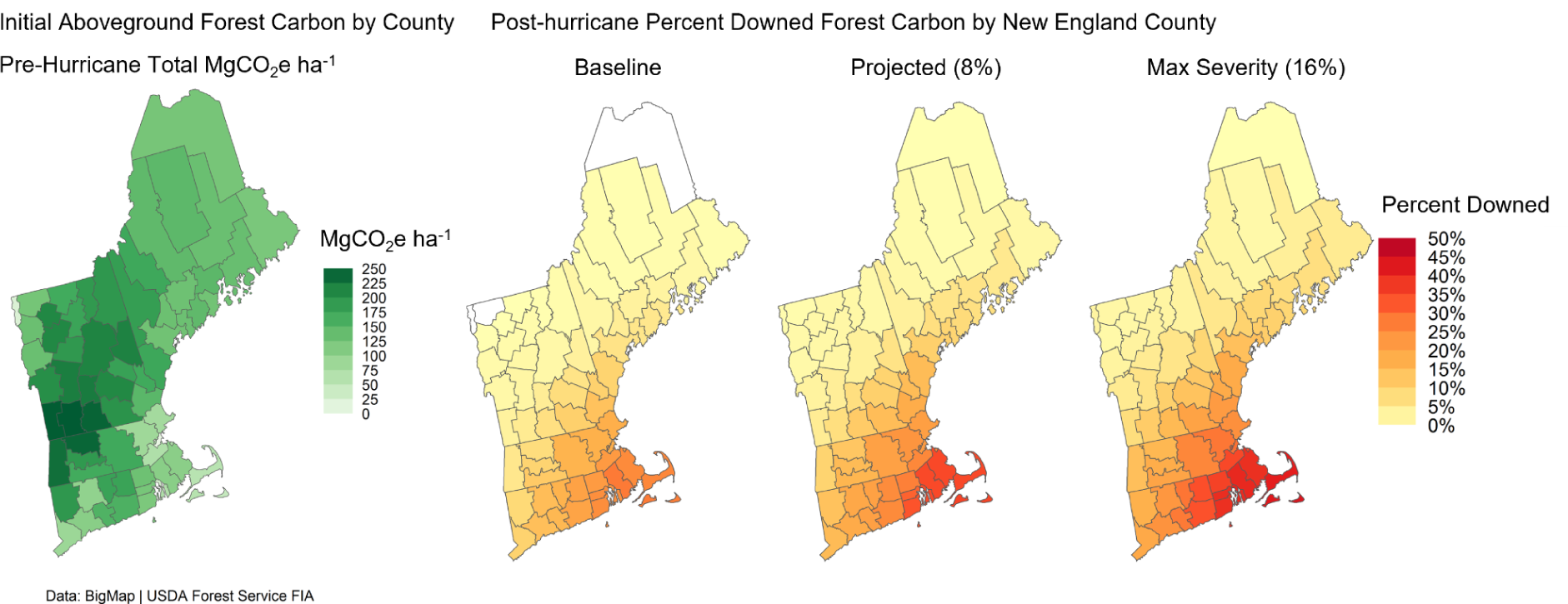
Initial (pre-hurricane) live aboveground forest carbon density in MgCO_2_e ha^-1^ (left) and the average percent of forest carbon downed by a hurricane (DFC/AFC*100) across the three scenarios (right) summarized by New England counties. The AFC values represent the carbon stored across our eight hardwood and softwood pools (Table 2), with dark green shades representing high forest carbon density and light green shades representing low forest carbon. Darker red and orange colors represent higher fractions of DFC, with lighter yellow shades represent lower percentages of DFC, and white represents zero DFC. Alternatively, Figure S1 shows the cumulative AFC and DFC values across New England counties in MMTCO_2_e.

**Table 4.** The total forested area (km^2^), proportion of total state area that is forested, and the aboveground forest carbon (AFC) density in Megagrams of carbon per hectare of state area (MgC ha^-1^).

| State | Forested Area (km<sup>2</sup>) | Percent Forested | AFC (MgC ha<sup>-1</sup>) |
| --- | --- | --- | --- |
| **Connecticut** | 7,200 | 56% | 70.1 |
| **Maine** | 70,467 | 83% | 45.8 |
| **Massachusetts** | 11,800 | 56% | 70.8 |
| **New Hampshire** | 19,326 | 80% | 68.4 |
| **Rhode Island** | 1,457 | 51% | 62 |
| **Vermont** | 18,509 | 74% | 68.3 |
| **New England** | **128,759** | **75%** | **56.3** |

Cumulatively across all of New England, there are 2,660 MMTCO_2_e AFC, with Rhode Island again having the lowest total AFC pool (33 MMTCO_2_e) and Maine having the largest (1,186 MMTCO_2_e; Table 5). Greenhouse gas flux data from the forest service show that the AFC pool in New England increases by 15.8 MMTCO_2_e on average annually, with New Hampshire and Vermont, being both heavily and densely forested (Table 4 & Figure 3; Walters et al. 2022), accounts for greater than half of the annual AFC flux (Table 5). The AFC flux was calculated using data from 2000-2020, a period with no major hurricane-induced DFC. The AFC and DFC values do not account for tree carbon in non-forested areas (25% of the landscape), or any carbon stored in plants that do not fall within our hardwood and softwood tree bins, such as shrubs, grasses, and forbs.

**Table 5.** Initial aboveground forest carbon (AFC) and annual AFC flux by state, along with total DFC (MMTCO_2_e) and percent downed per hurricane across the hurricane intensity scenarios.

|  | Initial Forest Conditions (MMTCO<sub>2</sub>e) |  | Baseline |  | Projected (8%) |  | Max Severity (16%) |  |
| --- | --- | --- | --- | --- | --- | --- | --- | --- |
|  | Annual AFC flux* | Total Initial AFC | DFC | % Down | DFC | % Down | DFC | % Down |
| Connecticut | -2.2 | 185 | 26 | 14.3% | 36 | 19.3% | 46 | 25.0% |
| Maine | -2.4 | 1,186 | 16 | 1.3% | 30 | 2.6% | 49 | 4.1% |
| Massachusetts | -2.9 | 307 | 42 | 13.6% | 57 | 18.7% | 71 | 23.2% |
| New Hampshire | -4 | 485 | 23 | 4.8% | 36 | 7.5% | 51 | 10.4% |
| Rhode Island | -0.2 | 33 | 8 | 23.5% | 10 | 30.0% | 13 | 37.6% |
| Vermont | -4.1 | 464 | 7 | 1.5% | 13 | 2.7% | 21 | 4.5% |
| **New England** | **-15.8** | **2,660** | **121** | **4.6%** | **182** | **6.9%** | **250** | **9.4%** |

\* Flux Data from Walters et al. (2022)

### 3.2 Extent of Hurricane Damage Across Hurricane Wind-intensity Scenarios

We modeled the 10 most impactful hurricanes of the 20th century under three wind intensity scenarios to estimate the damage that a hurricane would have on AFC under current forest conditions. The three scenarios are 1) baseline—reconstruction of hurricane extents and damage; 2) projected—8 percent increase in wind speeds from the baseline; and 3) maximum severity—16% increase in wind speeds. The average hurricane in the baseline scenario downs 4.6% (SD = 3%) of AFC, while hurricanes under the projected and max severity scenarios down 6.9% and 9.4% of AFC respectively (Table 5 & Figure 4). While the largest impacts occur in southern and coastal New England (Rhode Island, Connecticut, Massachusetts, and southern New Hampshire), increases in wind speeds lead to greater hurricane impacts both inland and northward (Figures 3 & S1), with the most drastic impacts being in the increased areas impacted by high-severity damage classes (EF2 & EF3).

**Figure 4.**
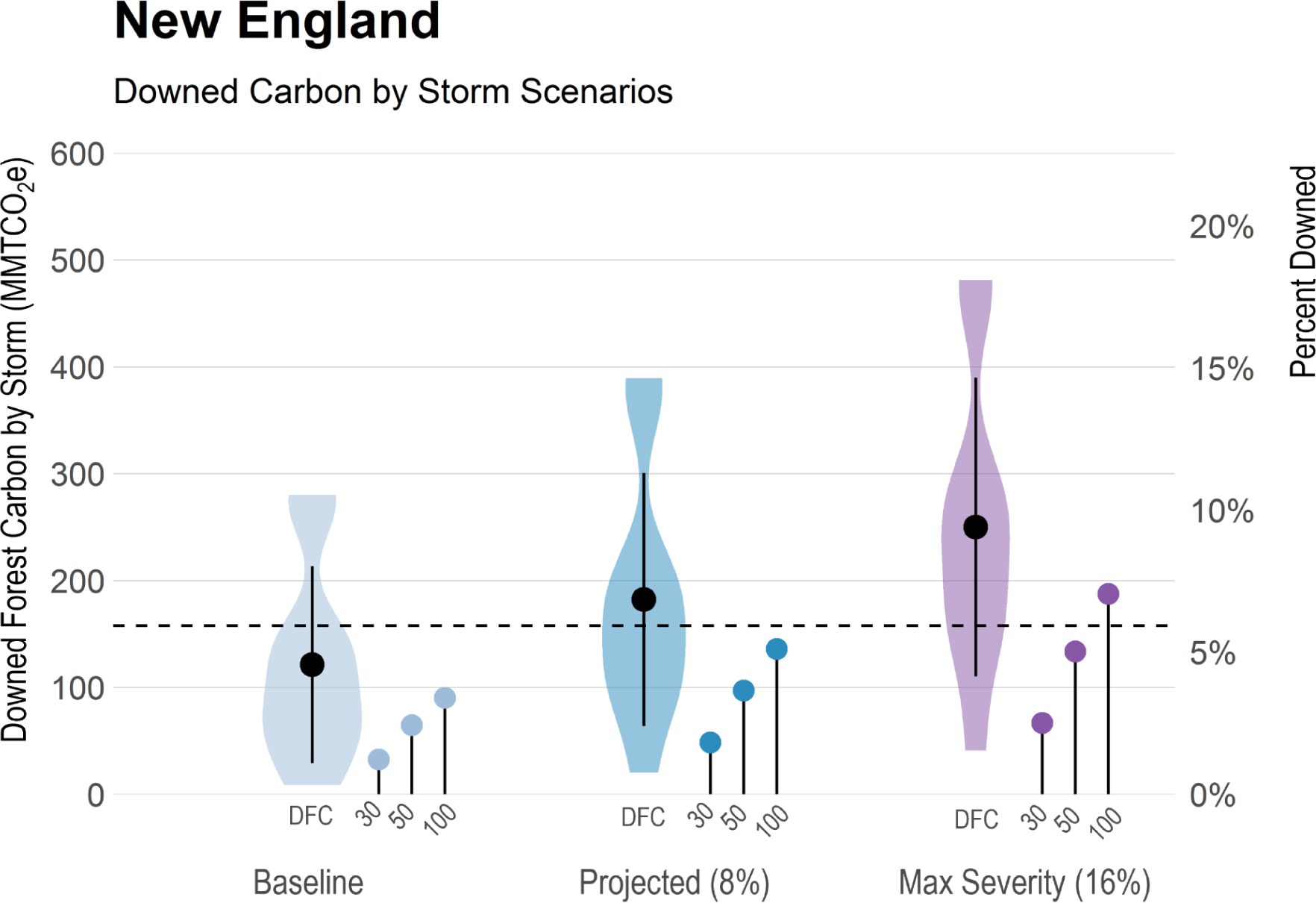
Average Downed Forest Carbon (DFC; MMTCO2e) for each hurricane across the hurricane intensity scenarios for all of New England. The black points represent the average DFC across the 10 hurricanes in each scenario (range represents the standard deviation) and the violins represent the distribution of DFC across hurricanes (Tables 2 & 5). The lollipop plots represent the cumulative net emissions from the average hurricane by scenario after 30, 50, and 100 years. The dashed line is the decadal carbon flux (absolute value–i.e., the decadal flux into the forest is -158 MMTCO2e) for New England forests (2000-2020) for reference (Walters et al. 2022). The secondary y-axis is the proportion of New England forest carbon downed by a storm (DFC/AFC*100).

On average, a hurricane from the baseline scenario impacts 4.3 million hectares (SD= 2.1 Mha) and results in 121 MMTCO_2_e of DFC (SD=92; Tables 6 & 5; Figures 4 & 5). The same hurricanes from the projected scenario (8% increase) impacts 6 million hectares and creates 182 MMTCO_2_e of DFC, an increase of 37.8% and 50.3% from the baseline scenario respectively. The maximum severity scenario (16% increase) storms have an average impact of 7.6 million hectares and creates 250 MMTCO_2_e of DFC, a DFC increase of 74.7% and 106.1% from the baseline scenario respectively.

**Figure 5.**
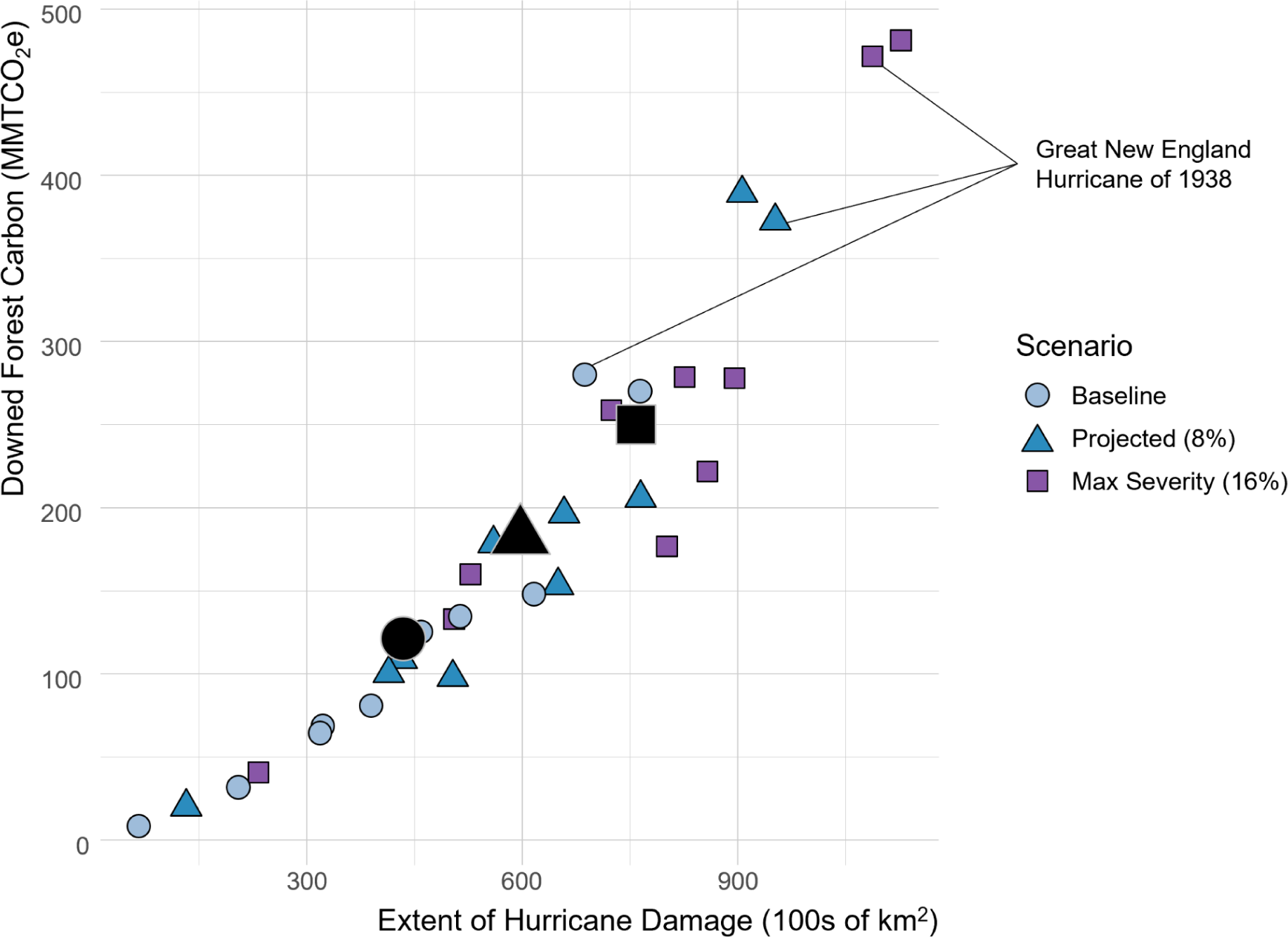
The extent of damage and downed forest carbon for the ten hurricanes we modeled across each of the three hurricane intensity scenarios. The points represent a single hurricane under each scenario: baseline (light blue circles), projected 8% wind speed increase (steel blue triangles), and the maximum severity scenario with a 16% wind speed increase (purple squares). The large black points represent the mean values for the 10 hurricanes under each scenario. The 1938 hurricane reconstructions are labeled to show the impact of increased wind speeds on an individual hurricane.

**Table 6.** The average impacted area (km^2^) by hurricane scenario across Enhanced Fujita (EF) class for New England states.

|  | Total Impacted Area |  | Damaged Area by EF Class (km<sup>2</sup>) |  |  |  |
| --- | --- | --- | --- | --- | --- | --- |
|  | km<sup>2</sup> | % | EF0 | EF1 | EF2 | EF3 |
| **Connecticut (12,933)** |  |  |  |  |  |  |
| Baseline | 8,548 | 66% | 3,971 | 3,548 | 935 | 94 |
| Projected | 10,057 | 78% | 3,969 | 4,061 | 1,556 | 471 |
| Max Severity | 10,969 | 85% | 3,444 | 3,953 | 2,666 | 905 |
| **Maine (84,903)** |  |  |  |  |  |  |
| Baseline | 7,513 | 9% | 6,787 | 726 | 0 | 0 |
| Projected | 14,611 | 17% | 12,970 | 1,513 | 128 | 0 |
| Max Severity | 23,300 | 27% | 19,850 | 3,130 | 320 | 0 |
| **Massachusetts (21,256)** |  |  |  |  |  |  |
| Baseline | 15,048 | 71% | 6,474 | 6,705 | 1,870 | 0 |
| Projected | 17,230 | 81% | 5,841 | 7,692 | 3,370 | 327 |
| Max Severity | 18,390 | 87% | 4,908 | 7,582 | 4,825 | 1,075 |
| **New Hampshire (24,039)** |  |  |  |  |  |  |
| Baseline | 7,453 | 31% | 5,524 | 1,915 | 14 | 0 |
| Projected | 10,709 | 45% | 7,451 | 2,915 | 342 | 0 |
| Max Severity | 13,398 | 56% | 8,047 | 4,437 | 913 | 0 |
| **Rhode Island (2,846)** |  |  |  |  |  |  |
| Baseline | 2,575 | 90% | 630 | 1,432 | 505 | 7 |
| Projected | 2,738 | 96% | 536 | 1,203 | 713 | 285 |
| Max Severity | 2,772 | 97% | 328 | 893 | 1,018 | 533 |
| **Vermont (24,903)** |  |  |  |  |  |  |
| Baseline | 2,263 | 9% | 1,985 | 278 | 0 | 0 |
| Projected | 4,369 | 18% | 3,721 | 639 | 8 | 0 |
| Max Severity | 6,999 | 28% | 5,837 | 1,061 | 101 | 0 |
| **New England (170,880)** |  |  |  |  |  |  |
| Baseline | 43,399 | 25% | 25,371 | 14,603 | 3,324 | 102 |
| Projected | 59,713 | 35% | 34,488 | 18,024 | 6,118 | 1,083 |
| Max Severity | 75,827 | 44% | 42,415 | 21,056 | 9,843 | 2,514 |

\*value in parentheses reflects the total area (km<sup>2</sup>) for each state/region

In the baseline scenario, 25% of the land area of New England is impacted by hurricanes on average. Most of the baseline scenario hurricane impacts are concentrated in southern New England, with less than 10% of the area of Vermont and Maine impacted by the average hurricane, and 90% of the damage in those two states are in the lowest damage class (i.e., EF0; Table 6), resulting in very little DFC (Table 3). Rhode Island, being the southernmost and coastal state, is the most affected by hurricane damage, with the average damaged area percent ranging from 90-97% across the hurricane scenarios (Table 6). As wind-intensities increase, 35% of New England is impacted under the projected scenario and 44% under the maximum-severity scenario (Table 6), with impacts shifting northwards and inland with increasing wind intensity (Figure 3). The largest impact across the scenarios came from the increase in the high-intensity EF3 damage from the baseline to the projected and maximum severity scenarios, with EF3 damage extent increasing by 1,066% and 2,475% respectively from the baseline (Table 6).

The ten hurricanes we modeled have a wide distribution in their extent and damage (Table 2), with the Great New England Hurricane (1938) and Hurricane Carol (1954) being the most damaging hurricanes of the 20th century (Figure 5—upper right points & Figure S2). For example, across the three wind severity scenarios, the 1938 hurricane affected 69-109 thousand km^2^ of New England land area and resulted in 280-472 MMTCO_2_e of DFC across the three hurricane scenarios (Table 2 & Figure S2). Alternatively, the weakest hurricane we included in our analysis, which occurred in 1962, impacted 7-23 thousand km^2^ of New England land area and resulted in only 8.6-41 MMTCO_2_e of DFC across the three hurricane scenarios (Table 2 & Figure S2).

### 3.3 Emissions Pathways of Downed Forest Carbon

We estimated the emissions pathways from downed wood in the forest following a hurricane disturbance. This included modeling estimates of the decay of DFC remaining in the forest, as well as using the harvested wood products to model the storage and emissions from the salvaged wood (estimated to be 25% of the total DFC pool). The salvaged wood is initially separated into various carbon pools, based on timber product ratios, and those pools are reconfigured through time based on the lifespans of timber products. Across scenarios, the fraction of DFC in the various pools are similar, given that the same timber product ratios are applied to the model, whereas differences in the pools are due to the composition of DFC (hardwood/softwood and tree height) and the overall magnitude of DFC (Figure S3).

Table 7 and Figure 6 show the carbon pools for key years following the disturbance (0, 30, 50, 100), while Figure S4 shows the continuous trajectory of the carbon pools for 100 years following the disturbance. Figure 7 displays the net emissions and the total stored and emitted DFC across the three scenarios. Immediately following a hurricane (year 0), most of the DFC is remaining in the forest (DFC_s_), with 4% of the DFC being emitted without energy capture (EWoEC), and 20% becoming timber products in use (PIU; Table 7, Figures 6 & S4). After 19 years, the DFC goes from a net store of carbon to a net emissions of carbon across all three scenarios (Figure 7), as the DFC_s_ remaining in the woods starts to decay and becomes emitted (DFC_e_), and the PIU carbon begins to be stored as solid waste (SWDS) or is emitted with or without energy capture (EEC & EWoEC; Figures 6 & S4). After 30 years, 64% of the total DFC has been emitted (Figure 4) and only the longest lived wood products remain as PIU. After 50 years, 77% of the initial DFC is emitted (Figure 4), with only a small fraction of the DFC remaining in the forest or in SWDS. After 100 years, 88% of the DFC is emitted (Figure 4), with ∼9% of the DFC in SWDS.

**Figure 6.**
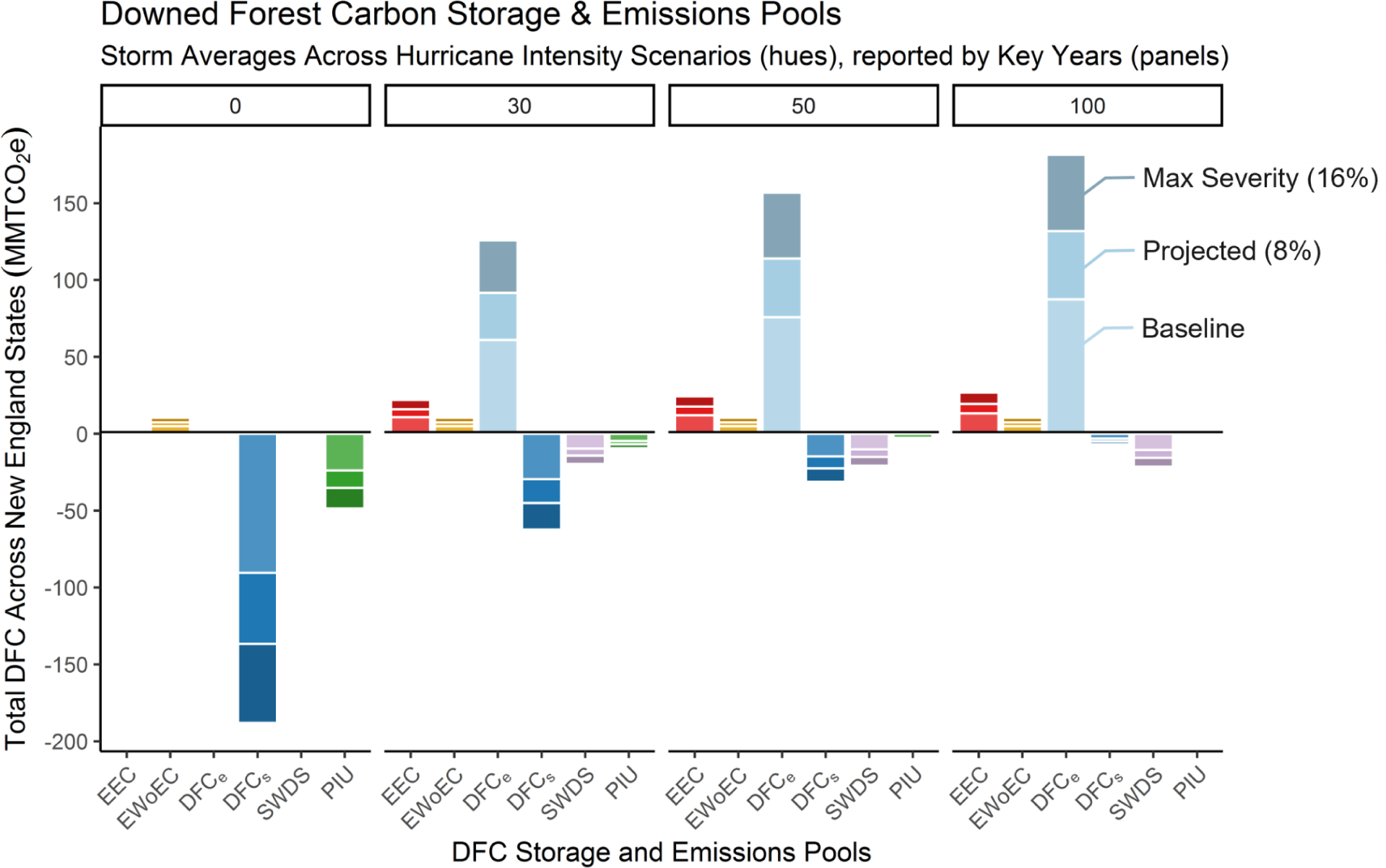
Storage and Emissions Pools of DFC based on the harvested wood products model across scenarios and through time. The y-axis is the average hurricane DFC for all of New England in MMTCO_2_e. Each panel represents certain years post hurricane disturbance. The bars represent the hurricane wind-intensity scenarios, with the baseline first in the lightest shade, followed by the projected scenario, and the maximum severity scenario in the darkest shade (see callout text in the last panel for example). The bars are cumulative, not stacked (i.e., the Projected value is the overall height of both the baseline and projected bars). The storage and emissions pools are: EEC—emitted with energy capture (fuelwood or burned onsite at mills for energy),EWoEC—emitted without energy capture (e.g., decay from SWDS), DFC_e_— decay/emissions from downed wood left in the forest. DFC_s_—downed wood remaining in the forest, SWDS—solid waste disposal sites such as dumps and landfills, and PIU—products in use. Figure S4 displays the continuous trajectory of DFC across the various storage and emissions pools for 100 years post-disturbance.

**Figure 7.**
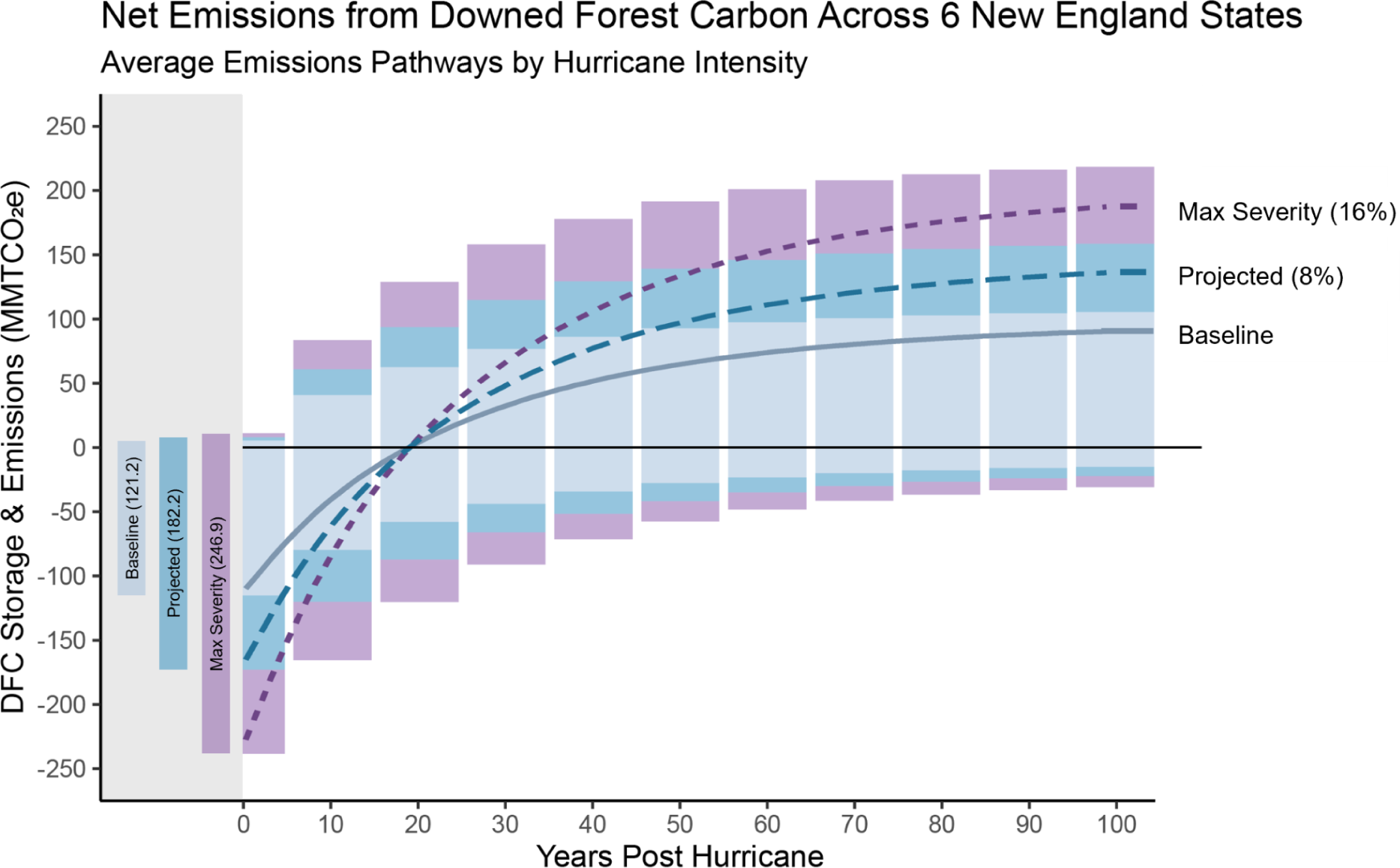
The net emissions from DFC across the baseline (light blue), projected (steel blue), and maximum severity (purple) scenarios following a hurricane according to the harvested wood products model. The gray region shows the total DFC pool, with the size of the pool in parentheses (MMTCO_2_e). The white region shows the trajectory of DFC as either storage (negative) or emissions (positive) pools across the scenarios through time. The bars are cumulative, not stacked. The lines depict the net emissions (storage + emissions).

**Table 7.**
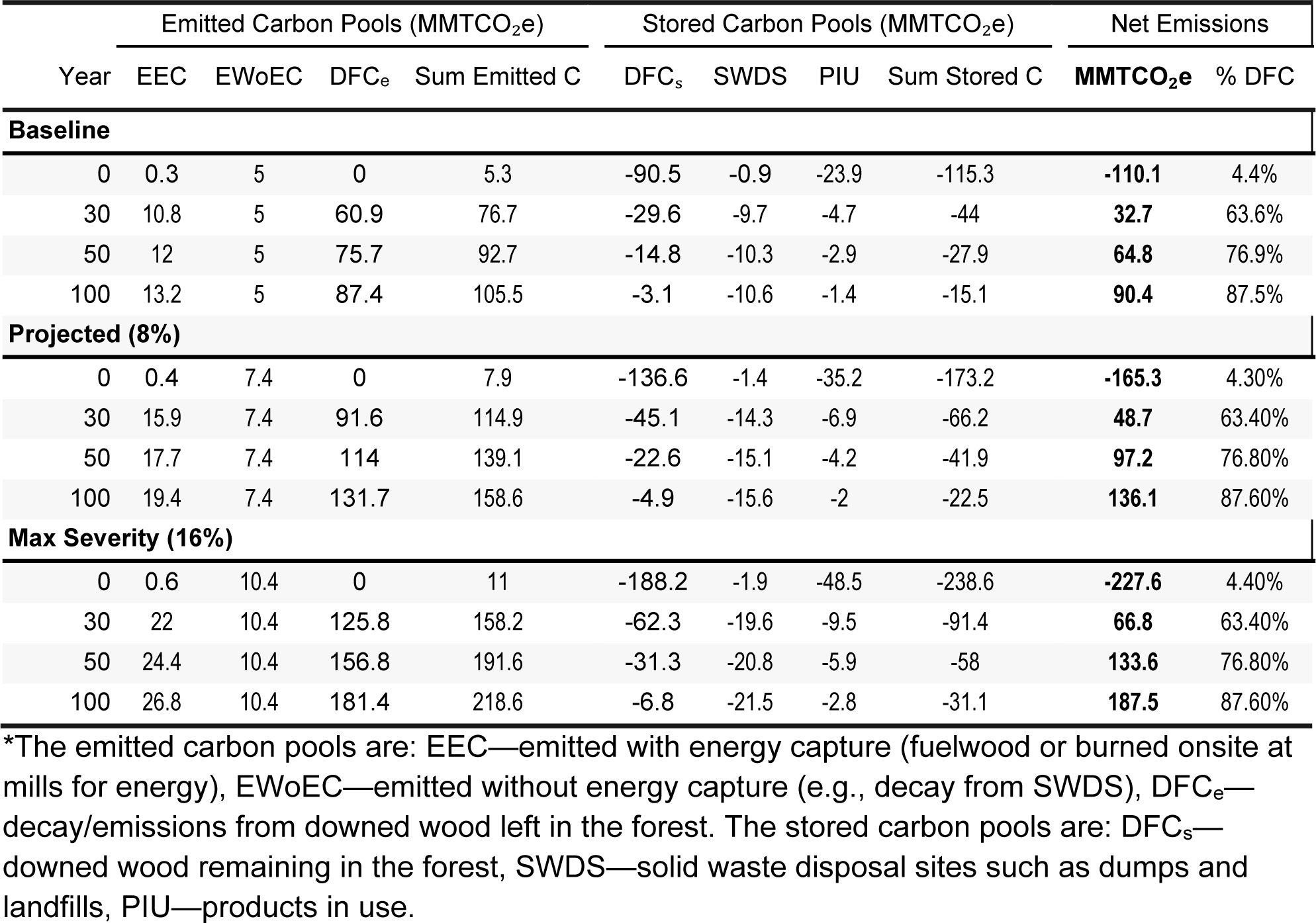
The trajectory of DFC carbon as either emitted or stored carbon following a hurricane across each scenario.

## 4.0 DISCUSSION

### 4.1 A single hurricane can emit decades-worth of carbon sequestration by New England’s forests

Across the hurricane scenarios, a single storm downs 121-250 MMTCO_2_e (4.6-9.4% of total aboveground forest carbon), the impact of which is much greater than the carbon sequestered annually across all of New England forests (15.8 MMTCO_2_e; Figure 4 & Table 5). Across the continental United States from 1980-1990, the CO_2_ released by hurricane damaged trees is equivalent to 9-18% of the forest carbon sink for that time period (Zeng et al. 2009). The majority of the impacts occur from a handful of large infrequent disturbances that have the capacity to alter landscapes and affect the net carbon flux (Foster et al. 1998; Zscheischler et al. 2014). For example, Hurricane Katrina in 2005 led to the mortality and damage of ∼320 million trees totalling 385 MMTCO_2_e, the equivalent of 50-140% of the net annual US forest tree carbon sink (Chambers et al. 2007). In our study, two out of the ten most impactful hurricanes of 20th century New England (The Great New England Hurricane of 1938 and Carol in 1954) accounted for 50% of the total aboveground forest carbon downed by hurricanes (Table 2 & Figure 5). Under the baseline scenario, without increased wind speeds, the impact of each of those two storms to the region is equivalent to roughly 18 years of the carbon sequestered by New England’s forests. The predicted warming of Atlantic basin sea surface temperatures as a result of climate change could strengthen hurricanes (Knutson et al. 2009), leading those two previous hurricanes to each negate 25-30 years of carbon sequestration across our projected and max severity scenarios.

### 4.2 Risks to New England Forests as a Nature-based Climate Solution (NCS): implications for forest offset schemes

Sequestering carbon in live forest biomass is widely considered as a premier NCS. With New England being one of the most heavily forested regions in the US, 75% forested by area (Table 4), containing 2,660 MMTCO_2_e of AFC, while also serving as an active carbon sink (Table 5), these forests are critical towards reaching our national and regional climate mitigation goals (Wayburn 2009; US Climate Change Science Program 2014; Thompson et al. 2020; Higgins et al. 2021). For example, Massachusetts is relying on the carbon sequestered in its natural and working lands (including forests) to sequester and “offset” up to 15% of its emissions as part of the Commonwealth’s Clean Energy and Climate Plan for 2050 (EEA 2022). New England forest landowners are also participating in the voluntary and compliance carbon markets, albeit only 6.7% of the forest carbon credits sold in California’s compliance market are in New England as of 2020 (Kaarakka et al. 2023), but participation in forest carbon offset schemes in the region is expected to increase, due to regulatory incentives and reduced participation barriers (Kerchner and Keeton 2015; Meyer et al. 2022; Pan et al. 2022). Current policies and carbon markets that rely on the land sector do not adequately account for the risks posed by hurricanes.

Using forest carbon as an offset for actualized emissions requires an adequate accounting of risks, to ensure that the forest carbon is additional, verifiable, and permanent (Roopsind et al. 2019; Badgley et al. 2022a). While there are considerable concerns regarding the actualized additionality and verifiability of forest offset credits, permanence is an exceptionally vulnerable aspect with regards to the viability of using temporary forest carbon pools to offset realized emissions (Pan et al. 2022; Haya et al. 2023). This is partly due to the fact that a change in forest land ownership could result in altered management practices that remove the “offsetted” carbon, but also because forests are vulnerable to ecological disturbances, such as fires, droughts, biological agents, and catastrophic risks (including severe-wind and precipitation), which could result in carbon losses (Hurteau et al. 2009; Ruseva et al. 2017). This is why many regulatory programs have created self-insurance programs to account for natural risks. For example, California’s cap-and-trade program, one of the largest regulatory markets for carbon offset credits, has created a buffer pool consisting of 8-12% of the credit to account for losses from natural risks (California Air Resources Board 2015). However, 95% of California’s buffer pool set-aside to mitigate fire risk (2-4% of all credits) has been depleted in less than 10% of the credits’ 100-year commitment (Badgley et al. 2022a).

The California buffer pool also includes a 3% discount for catastrophic risks like hurricanes, as well as other disturbance agents. Our results suggest that a single hurricane can down 4.6-9.4% of all AFC in New England, with southern New England forests expected to lose 13.6-37.6% of AFC, and northern New England forests 1.3-10.4% of AFC from any given storm (Table 5). Therefore, any single hurricane will likely deplete the buffer pool. This demonstrates that the risk to forest offsets from natural disturbances is significantly underestimated; therefore undermining the permanence and feasibility of using forest carbon to offset carbon emissions.

### 4.3 Stronger storms may lead to unprecedented impacts to northern and interior forests

Increases in hurricane wind speeds will likely lead to stronger and farther-reaching impacts. We found that the greatest increase in hurricane-induced forest carbon losses occurs due to the greater spatial extent of higher damage classes (EF2 & EF3). Meteorological predictions estimate that the frequency of category 4 and 5 hurricanes will double by the end of the 21st century, meaning that these higher impact storms could happen more frequently (Bender et al. 2010). Our hurricane reconstructions and projections found that, respectively, an 8% and 16% increase in wind speeds correspond to a 1,066% and 2,475% increase in the extent of EF3 level damage (where most trees are likely to succumb from wind-induced mortality). While most of the EF2 & EF3 damage is relegated to southern and coastal New England under the baseline scenario, stronger storms, as projected, will lead to unprecedented northward and inland shifts in high damage classes, affecting heavily forested regions in Western Massachusetts and northern New England. Extended land coverage from hurricanes has already been documented, as from the 1990s to 2000s there was a 63% increase in the length of hurricane-related storm tracks over U.S. land areas (Kasischke et al. 2013).

### 4.4 Disturbances agents differ in their forest carbon emissions consequences

Emissions from hurricane-induced downed forest carbon are not instantaneous, as it takes roughly 19 years for the carbon to transition from a net storage to a net emission, based on the decay rates of unsalvaged biomass and the lifespan of harvested timber products (Figures 7 & S4). Two thirds of the downed carbon is emitted after 30 years, 77% after 50 years, and 88% after 100 years (Table 7). Hurricanes differ substantially in their carbon consequences when compared to other disturbance types. For example, pyrogenic emissions from wildfires are emitted relatively instantaneously, and impact not just the live AFC, but can also combust necromass, litter, and soil carbon (Campbell et al. 2007; Chen et al. 2017). The residence time of hurricane-induced DFC left to decompose in the forest can be several decades, with an estimated necromass decay of 90% after 40 years across various temperate forests (Vrška et al. 2015; Khanina et al. 2023). The half-life of downed woody debris in eastern US forests is estimated to be 10 years for hardwoods and 20 years for softwoods (Russell et al. 2014).

Additionally, windthrown trees can often survive for a few years, demonstrated by the results of a simulated hurricane, where 90% of windthrown trees survive the first season, with 80% mortality within 6 years, and some trees even resprout or regrow after windthrow (Foster and Orwig 2006). Windthrow events also increase landscape heterogeneity by creating forest clearings and opportunities for the establishment of new species and by creating microsites through pit-and-mound topography from uprooting, and the influx of necromass can increase biodiversity and soil carbon (Franklin et al. 1987; Peterson and Pickett 1995; Carlton and Bazzaz 1998; Ulanova 2000).

Insect and disease outbreaks can also greatly alter the forest carbon balance directly and indirectly, differing substantially from hurricanes and wildfires, which are acute disturbances occurring over brief periods of time (Goetz et al. 2012; Kasischke et al. 2013). Insects/disease affect forests in various and complex ways: growth reducers (i.e., defoliation, herbivory, and disease) reduce tree productivity, whereas tree killers (i.e., bark beetles and pathogens) can lead to widespread mortality and reductions in forest carbon stocks (Hicke et al. 2012). These disturbances are difficult to estimate, because a variety of factors determine their impact: number of trees affected, density of targeted trees (insects/disease often impact specific species or groups), type of disturbance agent (growth reducers or tree killers), the duration of attack, and interactions with biotic and abiotic factors (such as other disturbance agents or climate Hicke et al. 2012). In New England, forests have been impacted by numerous biotic disturbance agents in recent decades, such as hemlock wooly adelgid which has decimated hemlock stands in southern New England, and Emerald Ash Borer which has rapidly spread leading to mortality within a few years of infestation (Orwig et al. 2008; Ignace et al. 2018; D’Amato et al. 2023). Many of the biotic agents in New England target specific species, leading to legacy shifts in forest composition; whereas windthrow indiscriminately impacts large and exposed trees, uprooting roughly 70% trees (compared to stem breakage), especially trees with unstable soils or root systems, or breaking trees that are more vulnerable to stem failure (Foster and Orwig 2006).

### 4.5 Emissions from downed forest carbon are influenced by salvage efficiency and timber product decisions

The emissions pathways and carbon consequences of hurricane-induced DFC is governed by three processes: 1) the decay rate of biomass left in the forest, 2) the salvage harvest efficiency, and 3) the half-lives of timber products from salvaged biomass. For our study, we assumed a 25% salvage rate based on historical trends and policy goals regarding disturbance responses, and limitations on timber processing, transportation, and storage capacity (Foster et al. 1997; Stanturf et al. 2007; Sanginés de Cárcer et al. 2021). In southern New England, the region most impacted by hurricanes, salvage capacity is incredibly low, as forestry has been declining steadily over the past several centuries. In the late 1930’s, Connecticut, Massachusetts, New Hampshire, and Vermont annually harvested about 500 million board feet of timber. The Great New England Hurricane of 1938 downed over 3 billion board feet, or about 70% of the merchantable timber in Central New England; therefore, the hurricane downed five years of timber harvests in just five hours. This spurred a massive response from the federal government and a previously declining forestry sector, as demonstrated by the rapidly increased number of active sawmills and storage sites for logs salvaged from the hurricane in the region, salvaging more than 1.5 billion board feet of lumber (Foster and Orwig 2006; Long 2016).

Would the forestry sector in New England respond at the scale necessary to salvage and process great quantities of timber following a disturbance? Northern New England has a larger forestry sector, but the largest impacts occur in Southern New England. The carbon emissions from salvaged wood products are dependent on the efficiency and products that the wood goes into. DFC used for biomass energy would be emitted rapidly, whereas salvaging timber for use in longer lived wood products would increase the length of time that the DFC is stored. Therefore, the ability to salvage greater quantities of DFC following a disturbance and to store that carbon in longer lived goods could decrease the carbon footprint of the disturbance. However, salvage harvests can also drastically alter biogeochemical cycles, leading to abrupt environmental and structural changes due to the disturbance caused by harvesting, whereas forests left to regenerate post disturbance have been capable of recovering rapidly with low to modest disruptions (Patric 1974; Bowden et al. 1993; Houlton et al. 2003; Foster and Orwig 2006).

### 4.6 Forest Recovery following hurricanes

We focused on the fate of New England forest carbon downed by a hurricane. Future research will examine the role of post-hurricane forest recovery on the carbon balance in the region. The impact that tropical cyclones have on the forest carbon balance in the U.S. is hotly debated. A synthesis of the forest carbon impacts from tropical cyclones across the continental U.S. from 1851-2000 found that tropical cyclones affect roughly 97 million trees per year, leading to an average carbon release of 92 MMTCO_2_e y^-1^ from DFC (Zeng et al. 2009). However, forest recovery following tropical cyclones has the potential of exceeding the carbon losses from downed trees, with the net annual flux of recovery potentially accounting for 17-36% of the US forest carbon sink (Fisk et al. 2013). The net carbon consequences of catastrophic wind events, such as hurricanes, on forest carbon remain unclear due to the difficulty of isolating the source-sink dynamics of the storms from other processes, and the impact that harvest and land-use decisions have on the carbon consequences of disturbances (Goetz et al. 2012; Williams et al. 2012). Future modeling will isolate the impacts that disturbances have on the net carbon balance of forests both immediately following the disturbance and throughout the recovery period.

## 5.0 CONCLUSIONS

Large infrequent disturbances, such as hurricanes, pose a major risk to the permanance of forest carbon stores. Our study of New England, one of the most forested regions of the U.S. and a significant carbon sink, demonstrates the impacts that hurricanes can have on forest carbon stocks and the risk of forests as nature-based climate solutions. Future research will investigate the recovery dynamics of post-disturbance forests and the long-term carbon balance of forested ecosystems. Here, we show that a single hurricane can emit decades-worth of carbon sequestered by forests, with New England hurricanes downing between 4.6-9.4% of all aboveground forest carbon in the region across our scenarios. Furthermore, we find that increases in hurricane wind speeds due to the projected warming of Atlantic basin sea surface temperatures could lead to unprecedented impacts both inland and northward into the heavily forested regions of northern New England.

## Acknowledgements

I would like to acknowledge everyone in the Thompson Lab at Harvard Forest who helped with the design and analysis of the study, especially Josh Plisinsiki and Lucy Lee. Thank you to David Foster for providing additional expertise and guidance. I would also like to thank the Harvard Forest LTER REU Program (NSF-DBI 1950364), and more broadly Harvard Forest and the NSF Funded LTER Program (NSF Grant No. DEB-LTER 18-32210) for supporting this research.

## Author Contributions Statement

**Emery Boose**: review & editing (equal), data curation (equal), software (equal), methodology (equal), conceptualization (equal). **Meghan Graham MacLean**: methodology (supporting), conceptualization (supporting), formal analysis (supporting), supervision (equal), software (equal), review & editing (equal). **Danelle Laflower**: review & editing (equal), conceptualization (supporting), methodology (supporting), formal analysis (supporting), visualization (equal). **Agustín León-Sáenz**: formal analysis (equal), data curation (equal), conceptualization (equal), methodology (equal), review & editing (supporting). **Taylor Lucey**: data curation (equal), formal analysis (equal), methodology (equal), software (equal), visualization (equal), original draft preparation (supporting), review & editing (equal). **Shersingh Joseph Tumber-Dávila**: conceptualization (equal), data curation (lead), formal analysis (lead), methodology (lead), visualization (lead), original draft preparation (lead), review & editing (equal). **Jonathan R. Thompson**: conceptualization (equal), project administration (lead), supervision (lead), original draft preparation (equal), review & editing (equal). **Barry T. Wilson**: software (equal), resources (equal), data curation (equal), review & editing (equal).

## Data Availability Statement

The data and scripts that support the findings of this study are openly available in the Environmental Data Initiative and Harvard Forest Archives at http://doi.org/[doi], reference number [reference number]. [Data and scripts have been submitted and are awaiting approval, the doi and reference number will be included prior to publication. Data citation will also be included once available.]

