## Supplemental Tables and Figures for "Hurricanes Pose Substantial Risk to New England Forest Carbon Stocks"

**Running Title:** Hurricanes Pose Risk to Forest Carbon

**Authors**:

Shersingh Joseph Tumber-Dávila
Harvard Forest, Harvard University, 324 N Main St, Petersham, MA 01366
Department of Environmental Studies, Dartmouth College, 39 College St, Hanover, NH 03755
0000-0001-7336-3943

Taylor Lucey
Department of Environmental Conservation, UMASS Amherst, 160 Holdsworth Way, Amherst, MA 01003
0000-0002-8880-9343

Emery Boose
Harvard Forest, Harvard University, 324 N Main St, Petersham, MA 01366
0000-0003-4820-0231

Danelle Laflower
Harvard Forest, Harvard University, 324 N Main St, Petersham, MA 01366

Agustín León-Sáenz
Harvard College, Harvard University, Massachusetts Hall, Cambridge, MA 02138
0009-0008-9459-2065

Barry T. Wilson
Northern Research Station, USDA Forest Service, 1992 Folwell Avenue, Saint Paul, MN 55108
0000-0002-1940-7682

Meghan Graham MacLean
Department of Environmental Conservation, UMASS Amherst, 160 Holdsworth Way, Amherst, MA 01003
0000-0002-5700-2168

Jonathan R. Thompson
Harvard Forest, Harvard University, 324 N Main St, Petersham, MA 01366
0000-0003-0512-1226

**Corresponding author**:
Shersingh Joseph Tumber-Dávila,, 978-756-6142

### Supplemental Figures

**Figure S1.** Initial (pre-hurricane) live aboveground forest carbon (AFC) in MMT$CO_2$e (left) and the average post-hurricane DFC across the three scenarios (right) summarized by New England counties. The AFC values represent the carbon stored across our eight hardwood and softwood pools (Table 2), with dark green shades representing high AFC stocks and light green shades representing low AFC stocks. Darker red and orange colors represent larger DFC pools with lighter yellow shades representing lower DFC, and white representing zero DFC. Figure 3 shows the AFC density per hectare and the fraction of AFC downed by a storm (DFC/AFC).

**Figure S2.** Maps of the ten individual hurricanes (rows) across the three hurricane intensity scenarios (columns) showing the hurricane path, enhanced fujita (EF) damage, and the proportion of downed forest carbon. The hurricane path is the black dashed line, while the contour lines represent the predicted EF damage based on the HURRECON/EXPOS models. The background raster is the percent of forest carbon downed by each storm, where gray is a non-forested pixel, beige is a forested pixel with zero damage, and the orange-red-purple colors represent increasing levels of downed aboveground forest carbon. The raster map is resampled from 30 to 90 meters, and represents the mean downed forest carbon for forested pixels.

**Figure S3.** Total initial aboveground forest carbon (AFC) in MMT$CO_2$e across the hardwood and softwood pools of various tree heights across New England States (green bars). The average downed forest carbon (DFC) from each hurricane across the three hurricane intensity scenarios: baseline (light blue), projected (steel blue), and maximum severity (purple).

**Figure S4.** The storage and emissions carbon pools of downed forest carbon following wind disturbances across the three hurricane intensity scenarios: Baseline (upper), Projected (middle), and Max Severity (lower). The carbon storage pools are products in use (PIU), solid waste disposal sites (SWDS), and remaining downed wood and residue left in the forest ($DFC_s$). The carbon emissions pools are emitted with energy capture (EEC), emitted without energy capture (EWoEC), and the decay of remaining DFC ($DFC_e$). The gray dashed line is the net carbon emissions (emitted + stored) from the DFC.

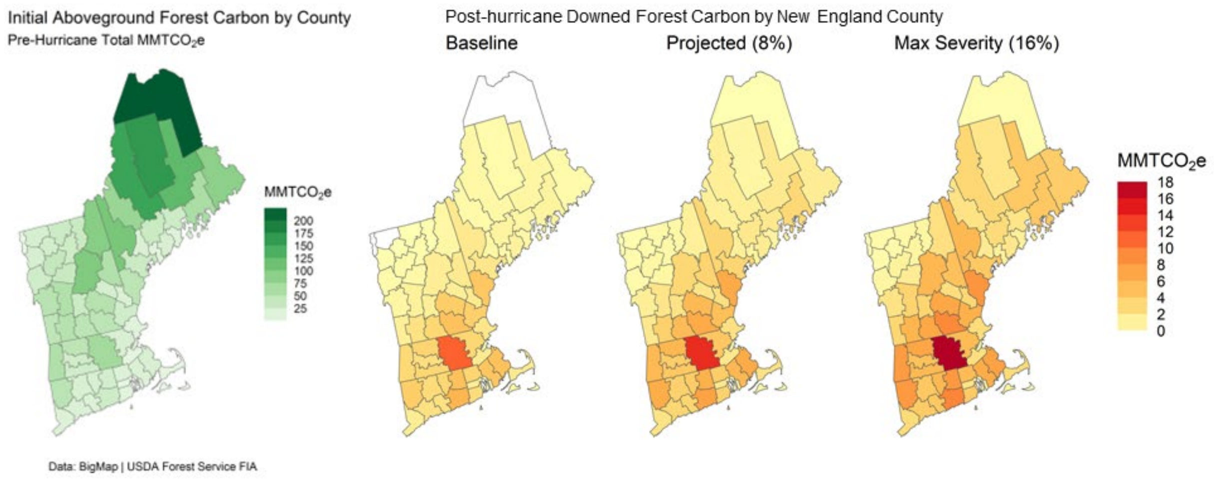


**Figure S1.** Initial (pre-hurricane) live aboveground forest carbon (AFC) in MMT$CO_2$e (left) and the average post-hurricane DFC across the three scenarios (right) summarized by New England counties. The AFC values represent the carbon stored across our eight hardwood and softwood pools (Table 2), with dark green shades representing high AFC stocks and light green shades representing low AFC stocks. Darker red and orange colors represent larger DFC pools with lighter yellow shades representing lower DFC, and white representing zero DFC. Figure 3 shows the AFC density per hectare and the fraction of AFC downed by a storm (DFC/AFC).

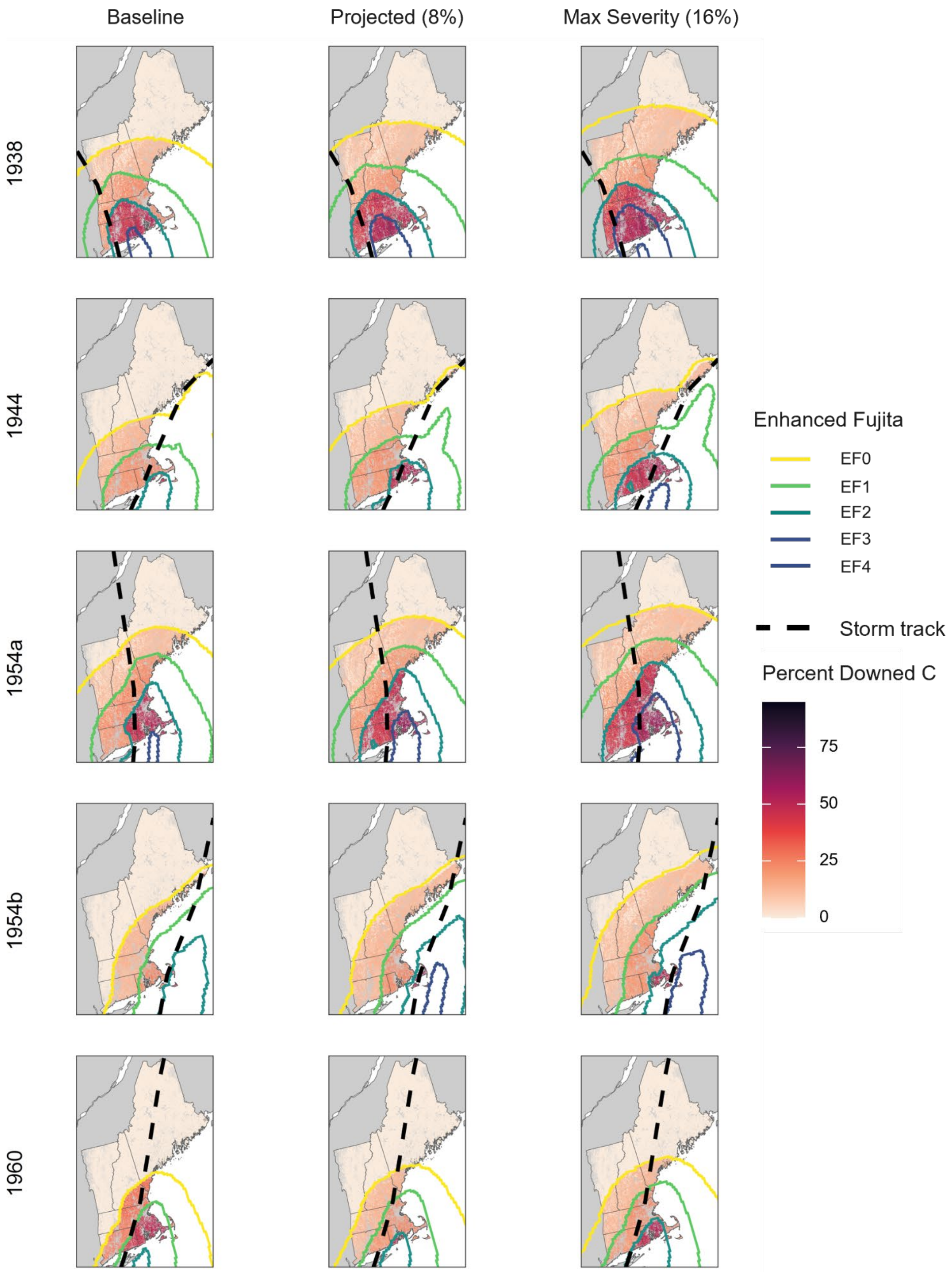

Baseline
Projected (8%)
Max Severity (16%)
1938
1944
1954a
1954b
1960
Enhanced Fujita
EF0
EF1
EF2
EF3
EF4
Storm track
Percent Downed C
75
50
25
0

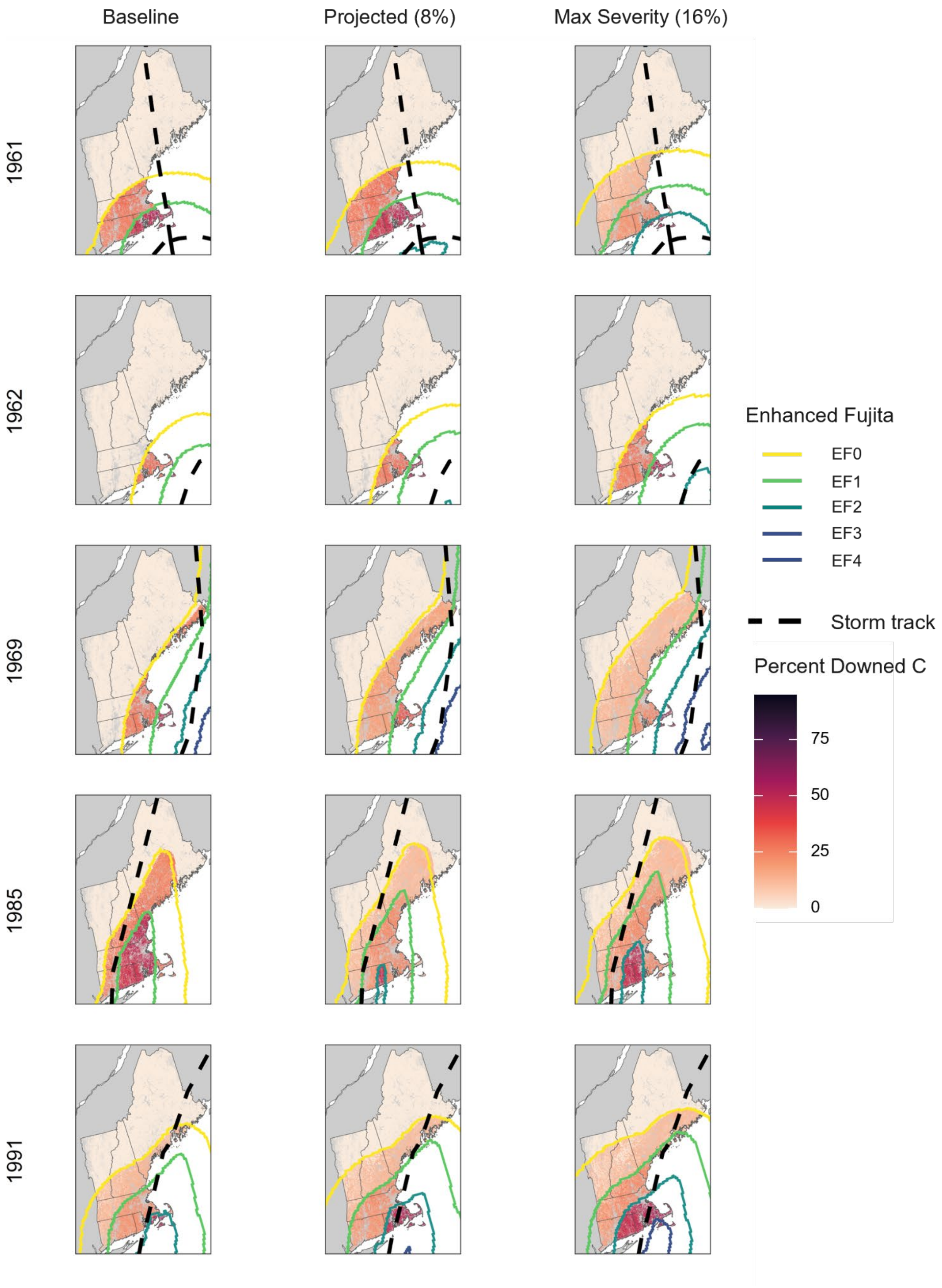
Baseline
Projected (8%)
Max Severity (16%)
1961
1962
1969
1985
1991
Enhanced Fujita
EF0
EF1
EF2
EF3
EF4
Storm track
Percent Downed C
75
50
25
0

**Figure S2.** Maps of the ten individual hurricanes (rows) across the three hurricane intensity scenarios (columns) showing the hurricane path, enhanced fujita (EF) damage, and the proportion of downed forest carbon. The hurricane path is the black dashed line, while the contour lines represent the predicted EF damage based on the HURRECON/EXPOS models. The background raster is the percent of forest carbon downed by each storm, where gray is a non-forested pixel, beige is a forested pixel with zero damage, and the orange-red-purple colors represent increasing levels of downed aboveground forest carbon. The raster map is resampled from 30 to 90 meters, and represents the mean downed forest carbon for forested pixels.

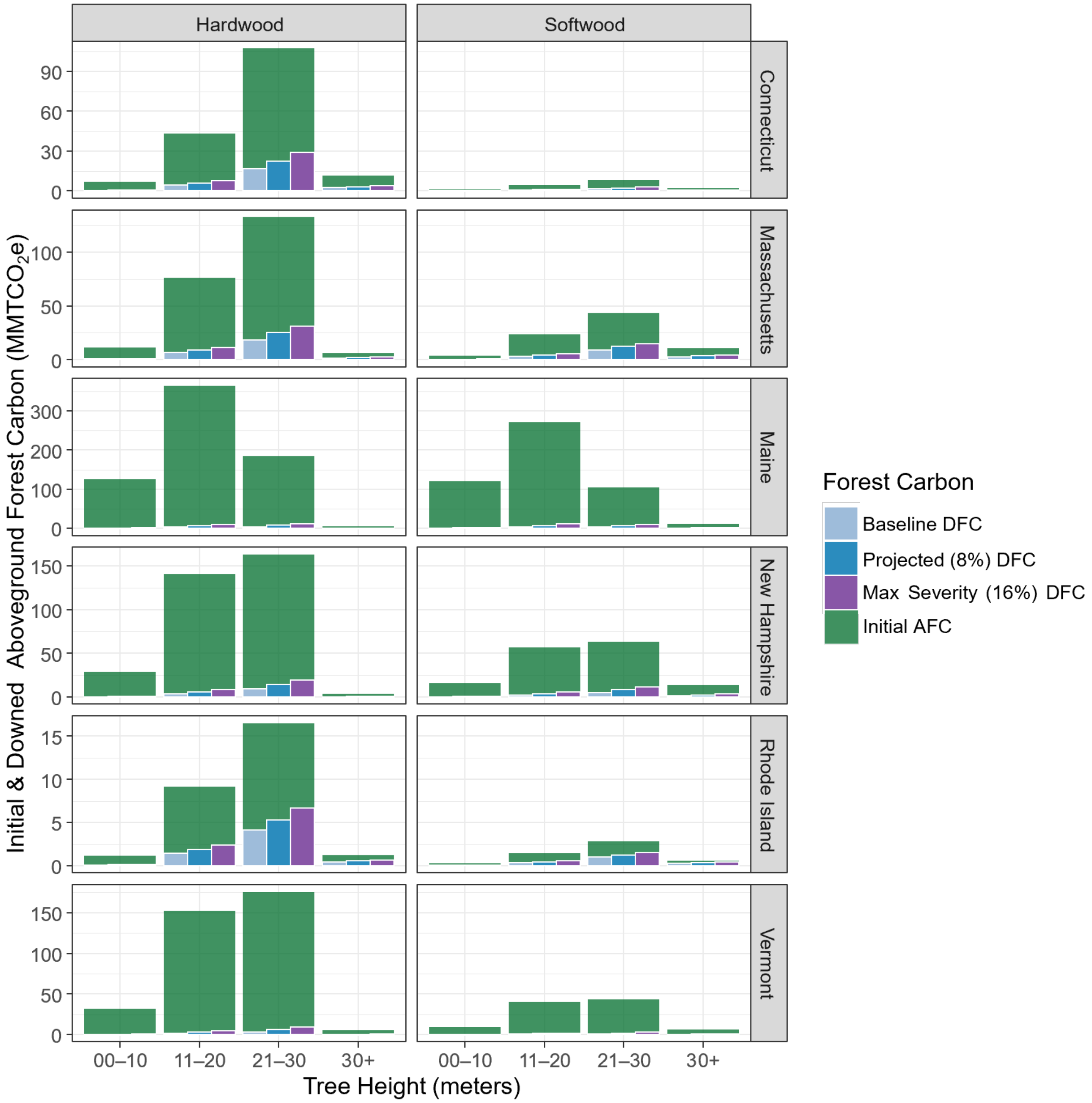


**Figure S3.** Total initial aboveground forest carbon (AFC) in MMTCO2e across the hardwood and softwood pools of various tree heights across New England States (green bars). The average downed forest carbon (DFC) from each hurricane across the three hurricane intensity scenarios: baseline (light blue), projected (steel blue), and maximum severity (purple).

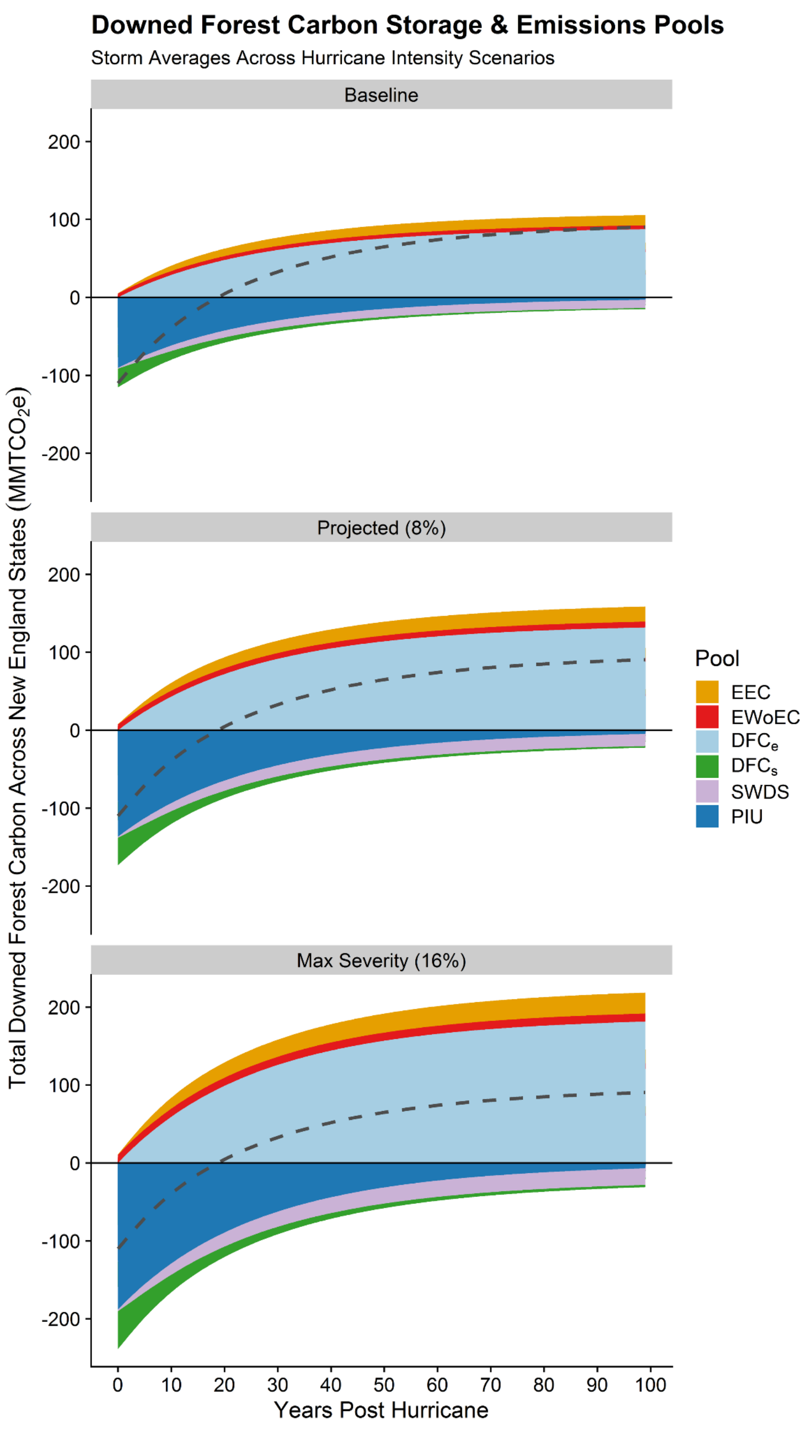


**Figure S4.** The storage and emissions carbon pools of downed forest carbon following wind disturbances across the three hurricane intensity scenarios: Baseline (upper), Projected (middle), and Max Severity (lower). The carbon storage pools are products in use (PIU), solid waste disposal sites (SWDS), and remaining downed wood and residue left in the forest ($DFC_s$). The carbon emissions pools are emitted with energy capture (EEC), emitted without energy capture (EWoEC), and the decay of remaining DFC ($DFC_e$). The gray dashed line is the net carbon emissions (emitted + stored) from the DFC.
